# The universal language network: A cross-linguistic investigation spanning 45 languages and 12 language families

**DOI:** 10.1101/2021.07.28.454040

**Authors:** Saima Malik-Moraleda, Dima Ayyash, Jeanne Gallée, Josef Affourtit, Malte Hoffmann, Zachary Mineroff, Olessia Jouravlev, Evelina Fedorenko

## Abstract

To understand the architecture of human language, it is critical to examine diverse languages; yet most cognitive neuroscience research has focused on a handful of primarily Indo-European languages. Here, we report an investigation of the fronto-temporo-parietal language network across 45 languages and establish the robustness to cross-linguistic variation of its topography and key functional properties, including left-lateralization, strong functional integration among its brain regions, and functional selectivity for language processing.

## Main Text

Approximately 7,000 languages are currently spoken and signed across the globe(Lewis, 2009). These are distributed across more than 100 language families—groups of languages that have descended from a common ancestral language, called the proto-language—which vary in size from 2 to over 1,500 languages. Certain properties of human languages have been argued to be universal, including their capacity for productivity (Chomsky, 1986) and communicative efficiency(Gibson et al., 2019). However, language is the only animal communication system that manifests in so many different forms(Evans & Levinson., 2009). The world’s languages exhibit striking diversity(Evans & Levinson, 2009), with differences spanning the sound inventories, the complexity of derivational and functional morphology, the ways in which the conceptual space is carved up into lexical categories, and the rules for how words can combine into phrases and sentences. To truly understand the nature of the cognitive and neural mechanisms that can handle the learning and processing of such diverse languages, we have to go beyond the limited set of languages used in most psycho- and neuro-linguistic studies(Bates et al., 1982; Bornkessel-Schlesewsky & Schlesewsky, 2016). This much needed step will also foster inclusion and representation in language research(Hudley et al., 2020).

Here, in a large-scale fMRI investigation, we evaluate the claim of language universality with respect to core features of its neural architecture. In the largest to date effort to sample many diverse languages, we tested native speakers of 45 languages across 12 language families (Afro-Asiatic, Austro-Asiatic, Austronesian, Dravidian, Indo-European, Japonic, Koreanic, Atlantic-Congo, Sino-Tibetan, Turkic, Uralic, and an isolate—Basque, which is effectively a one-language family). To our knowledge, about a third of these languages have never been investigated with functional brain imaging (or only probed in clinical contexts), no experimental paradigm has been tested with more than four languages at a time(Rueckl et al., 2015), and no attempts have been made to standardize tasks / language network definitions across languages, as needed to enable meaningful comparisons across studies (Supp. Table 1).

Using a powerful individual-subject analytic approach(Fedorenko et al., 2010), we examined the cross-linguistic generality of the following properties of the language network: i) topography (robust responses to language in the frontal, temporal, and parietal brain areas), ii) lateralization to the left hemisphere, iii) strong functional integration among the different regions of the network as assessed with inter-region functional correlations during naturalistic cognition, and iv) functional selectivity for language processing. All these properties have been previously shown to hold for English speakers. Because of their robustness at the individual-subject level(Mahowald & Fedorenko, 2016; Braga et al., 2020), and in order to test speakers of as many languages as possible, we adopted a ‘shallow’ sampling approach—testing a small number (n=2) of speakers for each language. The goal was not to evaluate any particular hypothesis/-es about cross-linguistic differences in the neural architecture of language processing (see discussion toward the end of the paper for examples), but rather to ask whether the core properties that have been attributed to the ‘language network’ based on data from English and a few other dominant languages extend to typologically diverse languages. Although we expected this to be the case, this demonstration—which can be construed as 45 conceptual replications (one for each language)—is an essential foundation for future systematic, in-depth, and finer-grained cross-linguistic comparisons. Another important goal was to develop robust tools for probing diverse languages in future neuroscientific investigations.

Each participant performed several tasks during the scanning session. First, they performed two language ‘localizer’ tasks: the English localizer based on the contrast between reading sentences and nonword sequences(Fedorenko et al., 2010) (all participants were fluent in English; Supp. Table 3), and a critical localizer task, where they listened to short passages from *Alice in Wonderland* in their native language, along with two control conditions (acoustically degraded versions of the native language passages where the linguistic content was not discernible and passages in an unfamiliar language). Second, they performed one or two non-linguistic tasks that were included to assess the functional selectivity of the language regions(Fedorenko et al., 2011) (a spatial working memory task, which everyone performed, and an arithmetic addition task, performed by 67 of the 86 participants). Finally, they performed two naturalistic cognition paradigms that were included to examine correlations in neural activity among the language regions, and between the language regions and regions of another network supporting high-level cognition: a ∼5 min naturalistic story listening task in the participant’s native language, and a 5 min resting state scan.

Consistent with prior investigations of a subset of these languages (e.g., Supp. Table 1), the activation landscape for the *Native-language>Degraded-language* contrast, which targets high-level language processing and activates the same set of brain areas as those activated by a more commonly used language localizer based on reading sentences versus nonword sequences(see Scott et al., 2017 for a direct comparison; also Supp. Figure 11), is remarkably consistent across languages and language families. The activations cover extensive portions of the lateral surfaces of left frontal, temporal, and parietal cortex (Figures 1, 2; see Supp. Figure 1, 2 for right hemisphere (RH) maps, and Supp. Figure 3 for volume-based maps). In the left-hemisphere language network (defined by the English localizer; see Supp. Figure 4 for evidence that similar results obtain in fROIs defined by the Alice localizer), across languages, the *Native-language* condition elicits a reliably greater response than both the *Degraded-language* condition (2.13 vs. 0.84 % BOLD signal change relative to the fixation baseline; t(44)=21.0, p<0.001) and the *Unfamiliar-language* condition (2.13 vs. 0.76; t(44)=21.0, p<0.001) (Figure 3a; see Supp. Figures 5-7 for data broken down by language, language family, and functional region of interest (fROI), respectively; see Supp. Table 2 for analyses with linear mixed effects models). Across languages, the effect sizes for the *Native-language>Degraded-language* and the *Native-Language>Unfamiliar-language* contrasts range from 0.49 to 2.49, and from 0.54 to 2.53, respectively; importantly, for these and all other measures, the inter-language variability is comparable to, or lower than, inter-individual variability (Supp. Figures 16-18).

**Figure 1.**
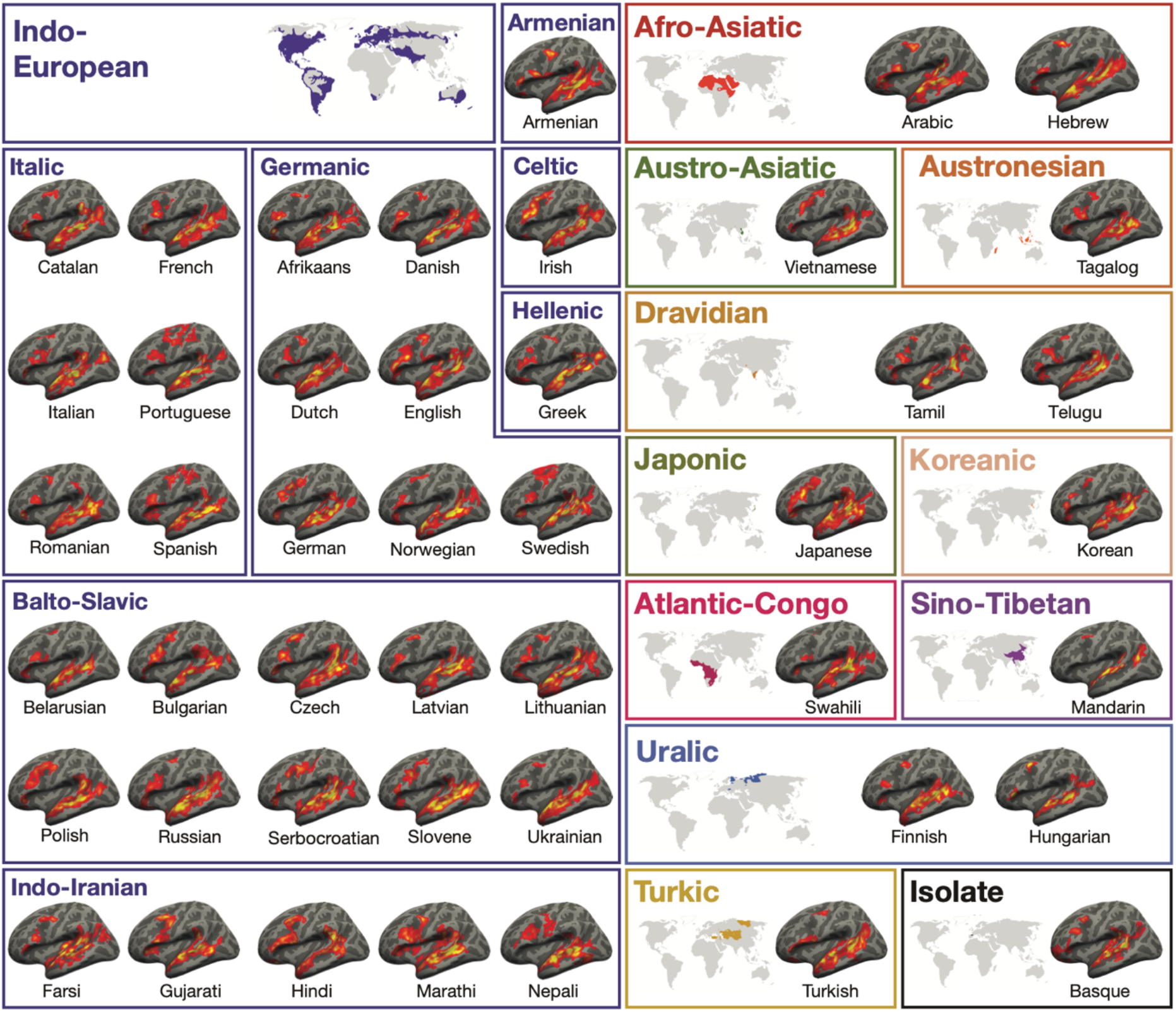
Activation maps for the Alice language localizer contrast (*Native-language>Degraded-language*) in the left hemisphere (LH) of a sample participant for each language (see Supp. Figure 1 for RH maps and details of the image generation procedure). The general topography of the language network in speakers of 45 languages is similar, and the variability observed is comparable to the variability that has been reported for the speakers of the same language (Mahowald & Fedorenko, 2016) (Supp. Figure 12). A significance map was generated for each participant by FreeSurfer(Dale et al., 1999); each map was smoothed using a Gaussian kernel of 4 mm full-width half-max and thresholded at the 70th percentile of the positive contrast for each participant. The surface overlays were rendered on the 80% inflated white-gray matter boundary of the fsaverage template using FreeView/FreeSurfer. Opaque red and yellow correspond to the 80th and 99th percentile of positive-contrast activation for each subject, respectively. (These maps were used solely for visualization; all the analyses were performed on the data analyzed in the volume (see Supp. Figure 3).)

**Figure 2.**
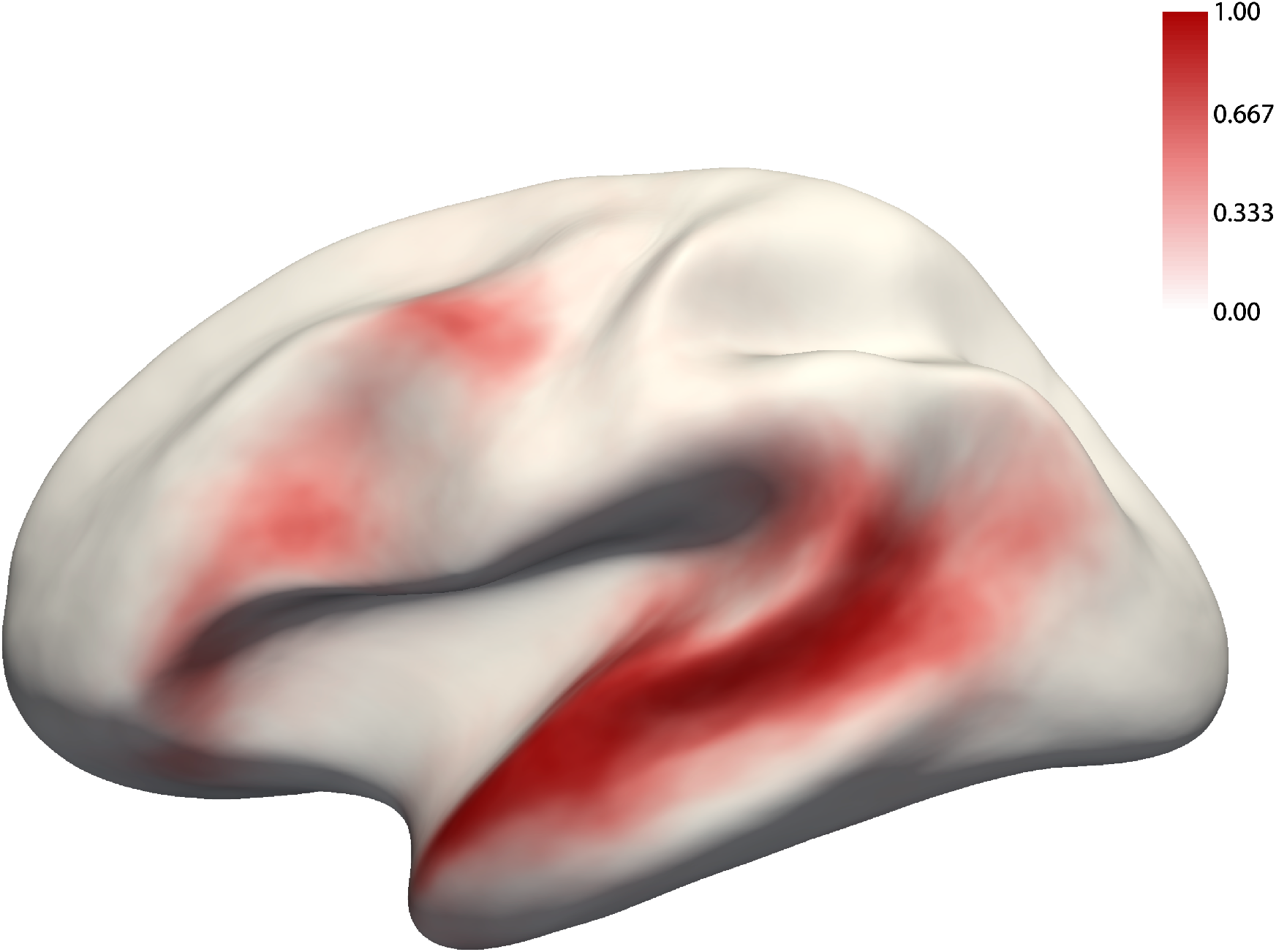
The probabilistic overlap map for the *Native-language>Degraded-language* contrast. This map was created by binarizing and overlaying the 86 participants’ individual maps (like those shown in Figure 1). The value in each vertex corresponds to the proportion of participants for whom that vertex belongs to the language network (see Supp. Figure 12 for a comparison between this probabilistic atlas vs. atlases based on native speakers of the same language).

**Figure 3.**
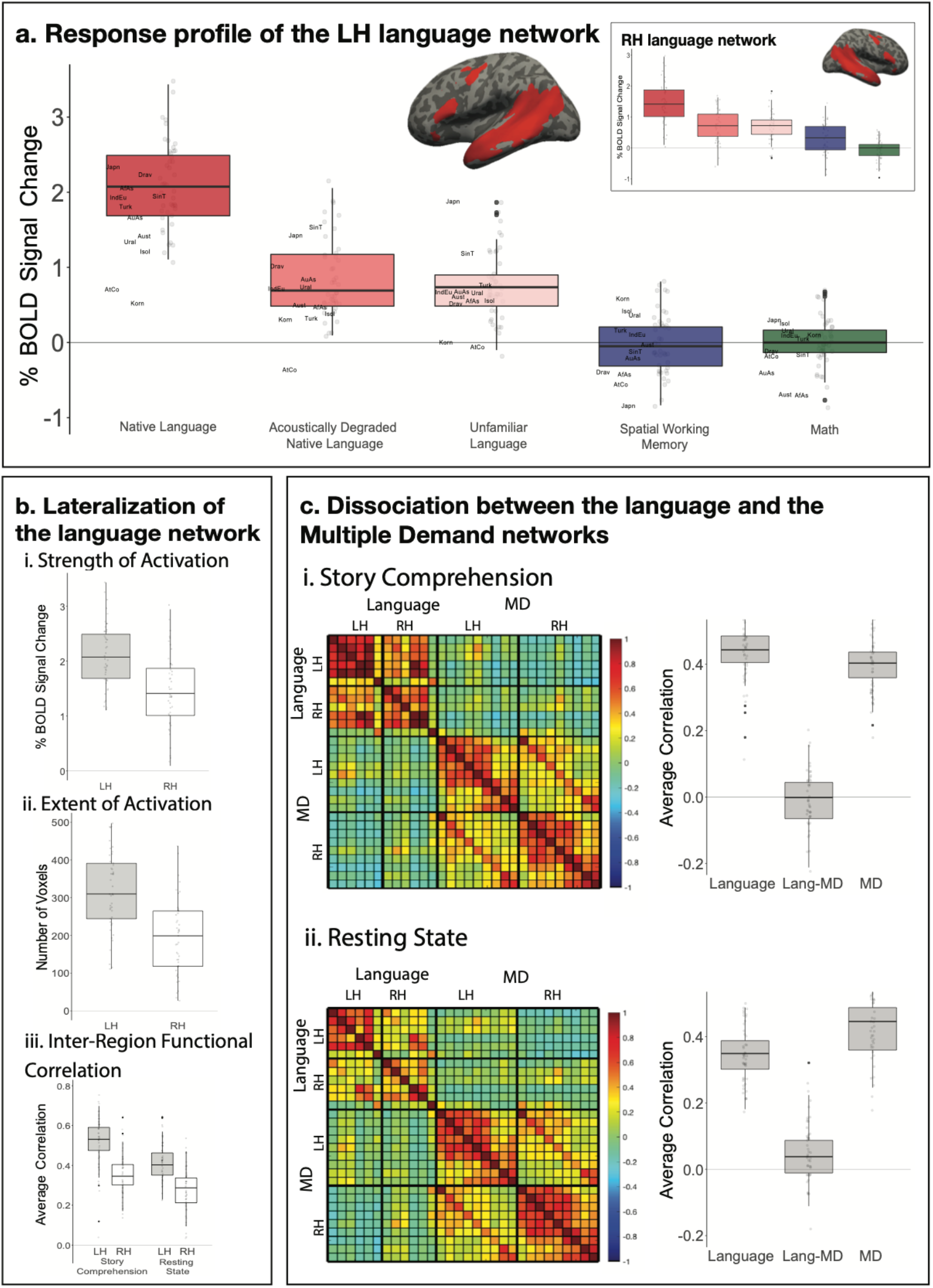
a) Percent BOLD signal change across the LH language functional ROIs (see inset for the RH language fROIs) for the three language conditions of the Alice localizer task (Native language, Acoustically degraded native language, and Unfamiliar language), the spatial working memory (WM) task, and the math task. The language fROIs show robust functional selectivity for language processing. Here and in the other panels, the dots correspond to languages (n=45), and the labels mark the averages for each language family (n=12; AfAs=Afro-Asiatic, AuAs=Austro-Asiatic, Aust=Austronesian, Drav=Dravidian, IndEu=Indo-European, Japn=Japonic, Korn=Koreanic, AtCo=Atlantic-Congo, SinT=Sino-Tibetan, Turk=Turkic, Ural=Uralic, Isol=Isolate). b) Three measures that reflect LH lateralization of the language network: i-strength of activation (effect sizes for the *Native-language>Degraded-language* contrast); ii-extent of activation (number of voxels within the union of the language parcels at a fixed threshold for the *Native-language>Degraded-language* contrast; a whole-brain version of this analysis yielded a similar result: t(44)=5.79, p<0.001); and iii-inter-region functional correlations during two naturalistic cognition paradigms (i-story comprehension in the participant’s native language; ii-resting state). The LH language network shows greater selectivity for language processing relative to a control condition, is more spatially extensive, and is more strongly functionally integrated than the RH language network. c) Inter-region functional correlations for the LH and RH language network and the Multiple Demand (MD) network during two naturalistic cognition paradigms (i-story comprehension in the participant’s native language; ii-resting state). The language and the MD networks are each strongly functionally integrated but are robustly dissociated from each other (pairs of fROIs straddling network boundaries show little/no correlated activity).

The *Native-language*>*Degraded-language* effect is stronger in the left hemisphere than the right hemisphere (2.13 vs. 1.47; t(44)=7.00, p<0.001), and more spatially extensive (318.2 vs. 203.5 voxels; t(44)=6.97, p<0.001; Figure 3b). Additionally, in line with prior data from English(Blank et al., 2014), the regions of the language network exhibit strong correlations in their activity during naturalistic cognition, with the average LH within-network correlation of r=0.52 during story comprehension and r=0.41 during rest, both reliably higher than zero (ts(44)>31.0, ps<0.001) and phase-shuffled baselines (ts(44)>10.0, ps<0.001; Figure 3c; see Supp. Figures 8 and 9 for data broken down by language). The correlations are stronger during story comprehension than rest (t(44)=-6.34, p<0.01). Further, as in prior work in English(Blank et al., 2014), and mirroring lateralization effects in the strength and extent of activation, the inter-region correlations in the LH language network are reliably stronger than those in the RH during both story comprehension (0.52 vs. 0.35; t(44)=8.00, p<0.001) and rest (0.41 vs. 0.28; t(44)=8.00, p<0.001; Figure 3c).

Finally, brain regions that support language processing have been shown to exhibit strong selectivity for language over many non-linguistic tasks, including executive function tasks, arithmetic processing, music perception, and action observation(Fedorenko et al., 2011; Fedorenko & Blank, 2020). This selectivity appears to be robustly present across speakers of diverse languages. Responses to the *Native-language* condition are significantly higher than those to the spatial working memory (WM) task (2.13 vs. −0.01; t(44)=20.7, p<0.001), and the math task (2.13 vs. 0.03; t(40)=21.5, p<0.001; Figure 3a, Supp. Figures 4-7). Furthermore, as in English(Blank et al., 2014), the language regions are robustly dissociated in their intrinsic fluctuation patterns from the regions of the bilateral domain-general multiple demand (MD) network implicated in executive functions(Duncan, 2010): within-network correlations are reliably greater than between-network correlations both during story comprehension (0.43 (language network, across the left and right hemisphere), 0.40 (MD network) vs. −0.01 (language-MD); ts(44)>23, ps<0.001), and during rest (0.34 (language, across hemispheres), 0.43 (MD) vs. −0.03 (language-MD), ts(44)>20, ps<0.001; Figure 3c, Supp. Figures 8, 9).

In summary, we have here established that key properties of the neural architecture of language hold across speakers of 45 diverse languages spanning 11 language families; and the variability observed across languages is comparable to, or lower than, the inter-individual variability among speakers of the same language(Mahowald & Fedorenko, 2016) (Supp. Figures 12, 16-18). Presumably, these features of the language network, including a) its *location* with respect to other—perceptual, cognitive, and motor—systems, b) *lateralization* to the left hemisphere (in most individuals), c) strong *functional integration* among the different components, and d) *selectivity* for linguistic processing, make it well-suited to support the broadly common features of languages, shaped by biological and cultural evolution.

In spite of their shared features, languages do exhibit remarkable variation(Evans & Levinson, 2009). How this variation relates to the neural implementation of linguistic computations remains a largely open question. By establishing broad cross-linguistic similarity in the language network’s properties and making publicly available the ‘localizer’ tasks (https://evlab.mit.edu/aliceloc) for 46 languages (to be continuously expanded over time), this work lays a critical foundation for future in-depth cross-linguistic comparisons along various dimensions of interest. In contrast to the shallow sampling approach adopted here (testing a small number of speakers across many languages), such investigations will require testing large numbers of speakers for each language / language family in question, while matching the groups carefully on all the factors that may affect neural responses to language. Such ‘deep’ sampling of each language / language family is necessary because cross-linguistic differences in the neural implementation of language processing are likely to be relatively subtle and they would need to exceed the (substantial) variability that characterizes speakers of the same language in order to be detected. Such investigations may also call for i) more fine-tuned/targeted paradigms (cf. the broad language contrast examined here), ii) multivariate analytic approaches, and iii) methods with high temporal resolution, like MEG or intracranial recordings(e.g., see Bornkessel-Schlesewsky et al., 2011; Bickel et al., 2015; Kemmerer, 2016 for past reports of cross-linguistic differences as measured with EEG). Regardless of the approach, the language localizer tasks enable narrowing in on the system of interest—the fronto-temporo-parietal network that selectively supports linguistic processing—thus yielding greater statistical power(Nieto-Castañón & Fedorenko, 2012), critical for detecting small effects, and interpretability, and leading to a robust and cumulative research enterprise.

What might hypotheses about cross-linguistic differences in neural implementation of language look like? Some examples include the following: i) languages with relatively strict word orders, compared to free-word-order languages, may exhibit a higher degree of left lateralization, given the purportedly greater role of the left hemisphere in auditory and motor sequencing abilities(Albert, 1972; Grafton et al., 2002; Poeppel, 2003), or stronger reliance on the dorsal stream, for similar reasons(Rauschecker, 2012; Bornkessel-Schlesewsky et al., 2015); ii) tonal languages may exhibit stronger anatomical and functional connections between auditory areas that process pitch(Norman-Haignere et al., 2013) and the higher-level language areas given the need to incorporate pitch information in interpreting word meanings(see Li et al., 2021 for evidence of a cross-linguistic difference in the lower-level speech perception cortex between speakers of a tonal vs. a non-tonal language); and iii) languages where utterances tend to underdetermine the meaning, like Riau Indonesian(Gil, 2013), may place greater demands on inferential processing to determine speaker intent and thus exhibit stronger reliance on brain areas that support such processes, like the right hemisphere language areas(Beeman, 1993) and/or the system that supports mental state attribution(Saxe&Kanwisher,2003).

Another class of hypotheses might come from the field of natural language processing (NLP). Recent advances in artificial intelligence have given rise to artificial neural network (ANN) models that achieve impressive performance on diverse language tasks(Radford et al., 2019; Devlin et al., 2018) and capture neural responses during language processing in the human brain (Schrimpf et al., 2021). Although, like cognitive neuroscience, NLP has also been dominated by investigations of English, there is growing awareness of the need to increase linguistic diversity in the training and evaluation of language models(Bender, 2009; Blasi et al., 2021), and some work has begun to probe cross-linguistic similarities and differences in the models’ learned representations(Chi et al., 2020; Papadimitriou et al., 2021). A promising future direction is to relate these cross-linguistic differences to neural differences observed during language processing across languages in an effort to illuminate how language implementation— in silico or in biological tissue—may depend on the properties of a particular language. More generally, because searching for cross-linguistic neural differences is a relatively new direction for language research(cf. Bornkessel-Schlesewsky & Schlesewsky, 2016), it will likely require a combination of top-down theorizing and bottom-up discovery. But no matter what discoveries about cross-linguistic differences in neural implementation lie ahead, the ability to reliably identify the language network in speakers of diverse languages opens the door to investigations of linguistic phenomena that are present in a small subset of the world’s languages, to paint a richer picture of the human language system.

Two limitations of the current investigation are worth noting. First, all participants were bilingual (fluent in English, in addition to their native language), which was difficult to avoid given that the research was carried out in the U.S. Some have argued that knowledge of two or more languages affects the neural architecture of one’s native language processing(Kovelman et al., 2008; Jouravlev et al., 2021). Importantly, however, i) this question remains controversial(Perani & Abutalebi, 2005; Costa & Sebastián-Gallés, 2014), and ii) we are not aware of any claims that learning a second language (L2) changes the properties of neural responses to the first language (L1) that we investigated here (i.e., their topography, lateralization, selectivity, or the strength of inter-regional correlations during naturalistic cognition). More generally, finding ‘pure’ monolingual speakers with no knowledge of other languages is challenging, especially in globalized societies, and is nearly impossible for some languages (e.g., Dutch, Galician, Kashmiri). The approach advocated here—where the language network is defined in each individual participant and individual-level neural markers are examined—allows for taking into account and explicitly modeling inter-individual variability in participants’ linguistic profiles (and along other dimensions), as will be important when evaluating specific hypotheses about cross-linguistic differences in future work, as discussed above. Another limitation is the over-representation of Indo-European languages (31 of the 45 languages). The analysis in Supp. Figure 6, which shows that the key statistics hold across language families, ameliorates this concern to some extent. Nevertheless, development of language localizers and collection of data for non-Indo-European languages remains a priority for the field. Our group will continue to develop and release the localizers for additional languages (https://evlab.mit.edu/aliceloc), and we hope other labs across the world will join this effort.

In conclusion, probing human language in all its diverse manifestations is critical for uncovering additional shared features, understanding the cognitive and neural basis of different solutions to similar communicative demands, characterizing the processing of unique/rare linguistic properties, and fostering diversity and inclusion in language sciences.

## Online Methods

### Participants

Ninety-one participants were recruited from MIT and the surrounding Boston community. Participants were recruited on the basis of their native language (the language acquired during the first few years of life; Supp. Table 3). All participants were proficient in English (Supp. Table 3). Data from 5 participants were excluded from the analyses due to excessive in-scanner motion or sleepiness. The final set included 86 participants (43 males) between the ages of 19 and 45 (M=27.52, SD=5.49; Supp. Table 4). All participants were right-handed, as determined by the Edinburgh Handedness Inventory(Oldfield, 1971) (n=83) or self-report (n=3), and had normal or corrected-to-normal vision. All participants gave informed written consent in accordance with the requirements of MIT’s Committee on the Use of Humans as Experimental Subjects (COUHES) and were paid for their participation.

Participants’ native languages spanned 12 language families (Afro-Asiatic, Austro-Asiatic, Austronesian, Dravidian, Indo-European, Japonic, Koreanic, Atlantic-Congo, Sino-Tibetan, Turkic, Uralic, Isolate (Basque)) and 45 languages (Supp. Table 3). We tested 2 native speakers per language (one male, one female) when possible; for 4 of the 45 languages (Tagalog, Telugu, Slovene, and Swahili), we were only able to test one native speaker.

### Experimental Design

Each participant completed i) a standard language localizer task in English(Fedorenko et al., 2010), ii) the critical language localizer in their native language, iii) one or two non-linguistic tasks that were included to assess the degree of functional selectivity of the language regions (a spatial working memory task, which everyone performed, and an arithmetic addition task, performed by 67 of the 86 participants), and iv) two naturalistic cognition paradigms that were included to examine correlations in neural activity among the language regions, and between the language regions and regions of another network supporting high-level cognition—the domain-general multiple demand (MD) network(Duncan, 2010) (a ∼5 min naturalistic story listening task in the participant’s native language, and a 5 min resting state scan). With the exception of two participants, everyone performed all the tasks in a single scanning session, which lasted approximately two hours. One participant performed the English localizer in a separate session, and another performed the spatial working memory task in a separate session. (We have previously established that individual activations are highly stable across scanning sessions(Mahowald & Fedorenko, 2016; see also Braga et al., 2020).)

#### Standard (English-based) language localizer

Participants passively read English sentences and lists of pronounceable nonwords in a blocked design. The *Sentences*>*Nonwords* contrast targets brain regions that support high-level linguistic processing, including lexico-semantic and combinatorial syntactic/semantic processes(Fedorenko et al., 2012; Blank et al., 2016). Each trial started with 100 ms pre-trial fixation, followed by a 12-word-long sentence or a list of 12 nonwords presented on the screen one word/nonword at a time at the rate of 450 ms per word/nonword. Then, a line drawing of a finger pressing a button appeared for 400 ms, and participants were instructed to press a button whenever they saw this icon, and finally a blank screen was shown for 100 ms, for a total trial duration of 6 s. The simple button-pressing task was included to help participants stay awake and focused. Each block consisted of 3 trials and lasted 18 s. Each run consisted of 16 experimental blocks (8 per condition), and five fixation blocks (14 s each), for a total duration of 358 s (5 min 58 s). Each participant performed two runs. Condition order was counterbalanced across runs. (We have previously established the robustness of the language localizer contrast to modality (written/auditory), materials, task, and variation in the experimental procedure(Fedorenko et al., 2010; Scott et al., 2017; Chen et al., 2021).)

#### Critical (native-language-based) language localizer

##### Materials

Translations of *Alice in Wonderland*(Carroll, 1865) were used to create the materials. We chose this text because it is one of the most translated works of fiction, with translations existing for at least 170 languages(Lindseth & Tannenbaum, 2015), and is suitable for both adults and children. Using the original (English) version, we first selected a set of 28 short passages (each passage took between 12 and 30 sec to read out loud). We also selected 3 longer passages (each passage took ∼5 min to read out loud) to be used in the naturalistic story listening task (see below). For each target language, we then recruited a native female speaker, who was asked to a) identify the corresponding passages in the relevant translation (to ensure that the content is similar across languages), b) familiarize themselves with the passages, and c) record the passages. In some languages, due to the liberal nature of the translations, the corresponding passages differed substantially in length from the original versions; in such cases, we adjusted the length by including or omitting sentences at the beginning and/or end of the passage so that the length roughly matched the original. We used female speakers because we wanted to ensure that the stimuli would be child-friendly (for future studies), and children tend to pay better attention to female voices(Wolff, 1963). Most speakers were paid for their help, aside from a few volunteers from the lab. Most of the recordings were conducted in a double-walled sound-attenuating booth (Industrial Acoustics). Materials for 3 of the languages (Hindi, Tamil, and Catalan) were recorded outside the U.S.; in such cases, recordings were done in a quiet room using a laptop’s internal microphone. We ensured that all recordings were fluent; if a speaker made a speech error, the relevant portion/passage were re-recorded. For each language, we selected 24 of the 28 short passages to be used in the experiment, based on length so that the target passages were as close to 18 s as possible. Finally, we created acoustically degraded versions of the target short passages following the procedure introduced in Scott et al.(Scott et al., 2017). In particular, for each language, the intact files were low-pass filtered at a pass-band frequency of 500 Hz. In addition, a noise track was created from each intact clip by randomizing 0.02-second-long periods. In order to produce variations in the volume of the noise, the noise track was multiplied by the amplitude of the intact clip’s signal over time. The noise track was then low-pass filtered at a pass-band frequency of 8,000 Hz and a stop frequency of 10,000 Hz in order to soften the highest frequencies. The noise track and the low-pass filtered copies of the intact files were then combined, and the level of noise was adjusted to a point that rendered the clips unintelligible. The resulting degraded clips sound like poor radio reception of speech, where the linguistic content is not discernible. In addition to the intact and degraded clips in their native language, we included a third condition: clips in an unfamiliar language (Tamil was used for 75 participants and Basque for the remaining 11 participants who had some exposure to Tamil during their lifetime). All the materials are available from the Fedorenko lab website: https://evlab.mit.edu/aliceloc (to be available upon publication; in the meantime, the materials are available from SMM upon request).

##### Procedure

For each language, the 24 items (intact-degraded pairs) were divided across two experimental lists so that each list contained only one version of an item, with 12 intact and 12 degraded trials. Any given participant was presented with the materials in one of these lists. Each list additionally contained 12 unfamiliar foreign language clips (as described above) chosen randomly from the set of 24. Participants passively listened to the materials in a long-event-related design, with the sound delivered through Sensimetrics earphones (model S14). The *Native-language* condition was expected to elicit stronger responses compared to both the *Degraded-language* condition(Scott et al., 2017) and the *Unfamiliar-language* condition(Chen et al., 2021) in the high-level language processing brain regions(Fedorenko et al., 2010). These language regions appear to support the processing of word meanings and combinatorial semantic/syntactic processes(Fedorenko et al., 2020), and these processes are not possible for the degraded or unfamiliar conditions. Each event consisted of a single passage and lasted 18 s (passages that were a little shorter than 18 s were padded with silence at the end, and passages that were a little longer than 18 s were trimmed down). We included a gradual volume fade-out at the end of each clip during the last 2 s, and the volume levels were normalized across the 36 clips (3 conditions * 12 clips each) in each set. The materials were divided across three runs, and each run consisted of 12 experimental events (4 per condition), and three fixation periods (12 s each), for a total duration of 252 s (4 min 12 s). Each participant performed three runs. Condition order was counterbalanced across runs.

#### Non-linguistic tasks

Both tasks were chosen based on prior studies of linguistic selectivity(Fedorenko et al., 2011). In the *spatial working memory* task, participants had to keep track of four (easy condition) or eight (hard condition) locations in a 3 x 4 grid(Fedorenko et al., 2011). In both conditions, participants performed a two-alternative forced-choice task at the end of each trial to indicate the set of locations that they just saw. Each trial lasted 8 s (see (Fedorenko et al., 2011) for the timing details). Each block consisted of 4 trials and lasted 32 s. Each run consisted of 12 experimental blocks (6 per condition), and 4 fixation blocks (16 s in duration each), for a total duration of 448 s (7 min 28 s). Each participant performed 2 runs. Condition order was counterbalanced across runs. Note that in the main analyses of this task and the math task, we averaged across the hard and easy conditions (but see Supp. Figure 14).

In the *arithmetic addition* task, participants had to solve a series of addition problems with smaller (easy condition) vs. larger (hard condition) numbers. In the easy condition, participants added two single-digit numbers. In the hard condition, participants added two numbers, one of which was double-digits. In both conditions, participants performed a two-alternative forced-choice task at the end of each trial to indicate the correct sum. Each trial lasted 3 s. Each block consisted of 5 trials and lasted 15 s. Each run consisted of 16 experimental blocks (8 per condition), and 5 fixation blocks (15 s in duration each), for a total duration of 315 s (5 min 15 s). Most participants performed 2 runs; 12 participants performed 1 run; 19 participants did not perform this task due to time limitations. Condition order was counterbalanced across runs when multiple runs were performed.

#### Naturalistic cognition paradigms

In the *story listening* paradigm, participants were asked to attentively listen to one of the long passages in their native language. The selected passage was 4 min 20 s long in English. Recordings in other languages were padded with silence or trimmed at the end, to equalize scan length across languages. The same 2 sec fade-out was applied to these clips, as to the shorter clips used in the critical experiment. In addition, each run included 12 s of silence at the beginning and end, for a total duration of 284 s (4 min 44 s). In the *resting state* paradigm, following Blank et al. (2014), participants were asked to close their eyes but to stay awake and let their mind wander for 5 minutes. The projector was turned off, and the lights were dimmed. **fMRI data acquisition.** Structural and functional data were collected on the whole-body 3 Tesla Siemens Trio scanner with a 32-channel head coil at the Athinoula A. Martinos Imaging Center at the McGovern Institute for Brain Research at MIT. T1-weighted structural images were collected in 179 sagittal slices with 1 mm isotropic voxels (TR = 2,530 ms, TE = 3.48 ms). Functional, blood oxygenation level dependent (BOLD) data were acquired using an EPI sequence (with a 90° flip angle and using GRAPPA with an acceleration factor of 2), with the following acquisition parameters: thirty-one 4mm thick near-axial slices, acquired in an interleaved order with a 10% distance factor; 2.1 mm x 2.1 mm in-plane resolution; field of view of 200mm in the phase encoding anterior to posterior (A >> P) direction; matrix size of 96 x 96; TR of 2,000 ms; and TE of 30 ms. Prospective acquisition correction(Thesen et al., 2000) was used to adjust the positions of the gradients based on the participant’s motion one TR back. The first 10 s of each run were excluded to allow for steady-state magnetization.

### fMRI data preprocessing and first-level analysis

fMRI data were analyzed using SPM12 and custom MATLAB scripts. Each subject’s data were motion corrected and then normalized into a common brain space (the Montreal Neurological Institute (MNI) template) and resampled into 2mm isotropic voxels. The data were then smoothed with a 4mm Gaussian filter and high-pass filtered at 128 s. For the language localizer task and the non-linguistic tasks, a standard mass univariate analysis was performed whereby a general linear model estimated the effect size of each condition in each experimental run. These effects were each modeled with a boxcar function (representing entire blocks/events) convolved with the canonical hemodynamic response function. The model also included first-order temporal derivatives of these effects, as well as nuisance regressors representing entire experimental runs, offline-estimated motion parameters, and outlier time points (i.e., time points where the scan-to-scan differences in global BOLD signal were above 5 standard deviations, or where the scan-to-scan motion was above 0.9 mm).

The naturalistic cognition paradigms (story listening and resting state) were preprocessed using the CONN toolbox(Whitfield-Gabrieli & Nieto-Castanon, 2012) with default parameters, unless stated otherwise. First, in order to remove noise resulting from signal fluctuations originating from non-neuronal sources (e.g., cardiac or respiratory activity), the first five BOLD signal time points extracted from the white matter and CSF were regressed out of each voxel’s time-course. White matter and CSF voxels were identified based on segmentation of the anatomical image(Behzadi et al., 2007). Second, the residual signal was band-pass filtered at 0.008-0.09 Hz to preserve only low-frequency signal fluctuations(Cordes et al., 2001).

To create aesthetically pleasing activation projection images for Figure 1, the data were additionally analyzed in FreeSurfer(Dale et al., 1999). Although all the analyses were performed on the data analyzed in the volume, these surface-based maps are available at OSF, along with the volume-analysis-based maps: https://osf.io/cw89s/.

### fROI definition and response estimation

For each participant, functional regions of interest (fROIs) were defined using the Group-constrained Subject-Specific (GSS) approach(Fedorenko et al., 2010), whereby a set of parcels or “search spaces” (i.e., brain areas within which most individuals in prior studies showed activity for the localizer contrast) is combined with each individual participant’s activation map for the same contrast.

To define the language fROIs, we used six parcels derived from a group-level representation of data for the *Sentences*>*Nonwords* contrast in 220 participants (Figure 3a). These parcels included three regions in the left frontal cortex: one in the inferior frontal gyrus (LIFG, 740 voxels; given that each fROI is 10% of the parcel, as described below, the fROI size is a tenth of the parcel size), one in its orbital part (LIFGorb, 370 voxels), and one in the middle frontal gyrus (LMFG, 460 voxels); and three regions in the left temporal and parietal cortex spanning the entire extent of the lateral temporal lobe and extending into the angular gyrus (LAntTemp, 1,620 voxels; LPostTemp, 2,940 voxels; and LAngG, 640 voxels). (We confirmed that parcels created based on the probabilistic overlap map for *Native-language>Degraded-language* contrast from the 86 participants in the current study are similar (Supp. Figure 10). We chose to use the ‘standard’ parcels for ease of comparison with past studies.) Individual fROIs were defined by selecting—within each parcel—the top 10% of most localizer-responsive voxels based on the *t-*values for the relevant contrast (*Sentences>Nonwords* for the English localizer). We then extracted the responses from these fROIs (averaging the responses across the voxels in each fROI) to each condition in the critical language localizer (native language intact, acoustically degraded native language, and unfamiliar language), and the non-linguistic tasks (averaging across the hard and easy conditions for each task). Statistical tests were then performed across languages on the percent BOLD signal change values extracted from the fROIs.

We used the English-based localizer to define the fROIs i) because we have previously observed(Chen et al., 2021) that the localizer for a language works well as long as a participant is proficient in that language (as was the case for our participants’ proficiency in English (Supp. Table 3); see also Supp. Figure 15 for evidence that our participants’ responses to the English localizer conditions were similar to those of native speakers), and ii) to facilitate comparisons with earlier studies(Fedorenko et al., 2011; Blank et al., 2014). However, in an alternative set of analyses (Supp. Figure 4), we used the *Native-language>Degraded-language* contrast from the critical language localizer to define the fROIs. In that case, to estimate the responses to the conditions of the critical language localizer, across-runs cross-validation(Nieto-Castañón & Fedorenko, 2012) was used to ensure independence(Kriegeskorte et al., 2010). The results were nearly identical to the ones based on the English localizer fROIs, suggesting that the two localizers pick out similar sets of voxels. Furthermore, for the two native speakers of English who participated in this study, the *Native-language>Degraded-language* contrast and the *Sentences>Nonwords* contrast are voxel-wise spatially correlated at 0.88 within the union of the language parcels (Fisher-transformed correlation(Silver & Dunlap, 1987); Supp. Figure 11). (Following a reviewer’s suggestion, we further explored the similarity of the activation maps for the *Native-language>Degraded-language* and *Native-language>Unfamiliar-language* contrasts in the Alice localizer. These maps were similar: across the 86 participants, the average Fisher-transformed voxel-wise spatial correlation within the union of the language parcels was 0.66 (SD = 0.40; see Supp. Figure 13 for sample individual map pairs), and the magnitudes of these effects did not differ statistically (t(44)=1.15, p=0.26). These results suggest that either contrast can be used to localize language-responsive cortex—along with the more traditional *Sentences>Nonwords* contrast—although we note that, among the two auditory contrasts, we have more and stronger evidence that the *Native-language>Degraded-language* works robustly and elicits similar responses to the *Sentences>Nonwords* contrast.)

In addition to the magnitudes of response, we estimated the degree of language lateralization in the native language localizer based on the extent of activation in the left vs. right hemisphere. To do so, for each language tested, in each participant, we calculated the number of voxels activated for the *Native-language>Degraded-language* contrast (at the *p*<0.001 whole-brain uncorrected threshold) within the union of the six language parcels in the left hemisphere, and within the union of the homotopic parcels in the right hemisphere(Mahowald & Fedorenko, 2016), as shown in Figure 2b. Statistical tests were then performed across languages on the voxel count values. (We additionally performed a similar analysis considering the voxels across the brain(Seghier, 2008).)

Finally, we calculated inter-regional functional correlations during each of the naturalistic cognition paradigms. For these analyses, in addition to the language fROIs, we examined a set of fROIs in another large-scale brain network that supports high-level cognition: the domain-general multiple demand (MD) network(Duncan, 2010, 2013), which has been implicated in executive functions, like attention, working memory, and cognitive control. This was done in order to examine the degree to which the language regions are functionally dissociated from these domain-general MD regions during rich naturalistic cognition, as has been shown to be the case for native English speakers(Blank et al., 2014; Paunov et al., 2019). To define the MD fROIs, following(Fedorenko et al., 2013; Blank et al., 2014), we used anatomical parcels(Tzourio-Mazoyer et al., 2002) that correspond to brain regions linked to MD activity in prior work. These parcels included regions in the opercular IFG, MFG, including its orbital part, insular cortex, precentral gyrus, supplementary and presupplementary motor area, inferior and superior parietal cortex, and anterior cingulate cortex, for a total of 18 regions (9 per hemisphere). Individual MD fROIs were defined by selecting—within each parcel—the top 10% of most localizer-responsive voxels based on the *t-*values for the *Hard>Easy* contrast for the spatial working memory task(Blank et al., 2014) (see Supp. Figure 14 for an analysis showing that this effect is highly robust in the MD fROIs, as estimated using across-runs cross-validation, as expected based on prior work).

For each subject, we averaged the BOLD signal time-course across all voxels in each language and MD fROI. We then averaged the time-courses in each fROI across participants for each language where two participants were tested. For each language, we computed Pearson’s moment correlation coefficient between the time-courses for each pair of fROIs. These correlations were Fisher-transformed to improve normality and decrease biases in averaging(Silver & Dunlap, 1987). We then compared the average correlation for each language a) within the language network (the average of all 66 pairwise correlations among the 12 language fROIs), b) within the MD network (the average of all 190 pairwise correlations among the 20 MD fROIs), and c) between language and MD fROIs (the average of 240 pairwise correlations between the language fROIs and the MD fROIs). For the language network, we also computed the within-network correlations for the left and right hemisphere separately, to examine lateralization effects. All the statistical comparisons were performed across languages. The fROI-to-fROI correlations are visualized in two matrices, one for each naturalistic cognition paradigm (Figure 3c).

### Data availability

The data that support the findings of this study are available at: https://osf.io/cw89s.

### Code availability

The code used to analyze the data in this study are available at: https://osf.io/cw89s.

## Acknowledgments

We thank i) Zoya Fan, Jorge Vera-Rebollar, Frankie Frank, Annemarie Verkerk, Celeste Kidd, and Ming Xiang for help with locating the texts of *Alice in Wonderland* in different languages; ii) Zoya Fan, Frankie Frank, and Jorge Vera-Rebollar for help with finding and recording the speakers; iii) Idan Blank, Alex Paunov, and Ben Lipkin for help with some of the analyses; iv) Josh McDermott for letting us use the sound booths in his lab for the recordings; v) Jin Wu, Niharika Jhingan, and Ben Lipkin for creating a website for disseminating the localizer materials and script; vi) Martin Lewis for allowing us to use the linguistic family maps from the GeoCurrents website; vii) Barbara Alonso Cabrera for help with figures; viii) EvLab and TedLab members and collaborators, and the audiences at the Neuroscience of Language Conference at NYU-AD (2019), and at the virtual Cognitive Neuroscience Society conference (2020) for helpful feedback, and Ted Gibson, Damián Blasi, and three anonymous reviewers for comments on earlier drafts of the manuscript; ix) Doug Greve and Bruce Fischl for their help with the FreeSurfer analyses; and x) our participants. The authors would also like to acknowledge the Athinoula A. Martinos Imaging Center at the McGovern Institute for Brain Research at MIT, and the support team (Steven Shannon and Atsushi Takahashi). S.M.-M. was supported by la Caixa Fellowship LCF/BQ/AA17/11610043. E.F. was supported by NIH awards R00-HD057522, R01-DC016607, and R01-DC-NIDCD and funds from the Brain and Cognitive Sciences Department, the McGovern Institute for Brain Research, and the Simons Center for the Social Brain.

## Author contributions

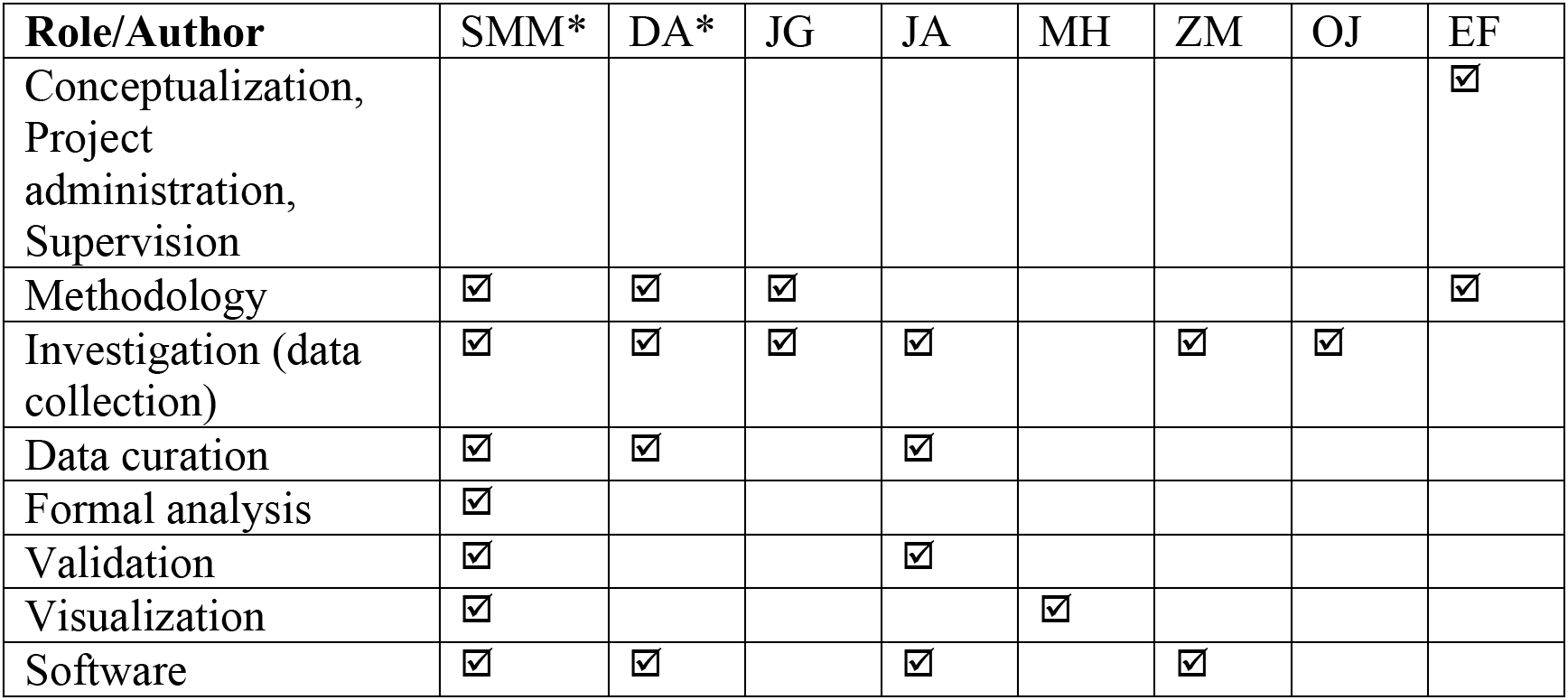
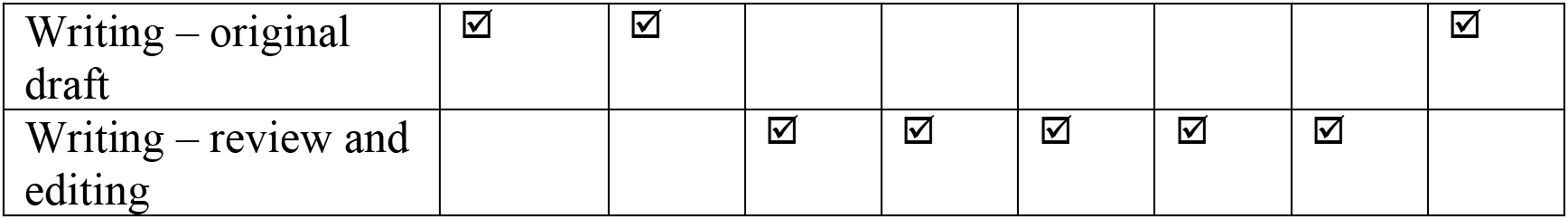

## Supplementary Information

### Supplementary Figures

**Supplementary Figure 1:**
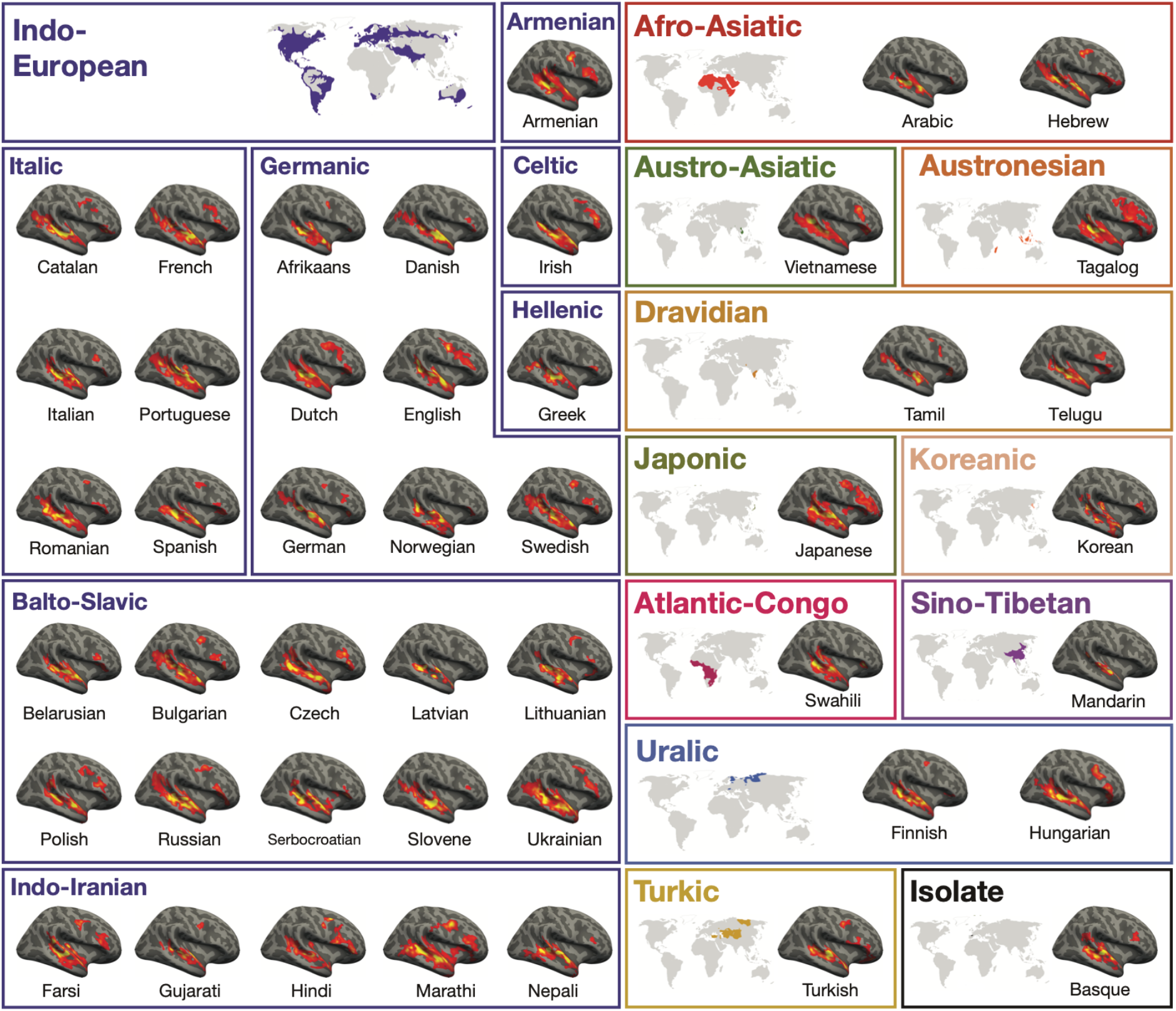
Activation maps for the Alice language localizer contrast (*Native-language>Degraded-language*) in the right hemisphere of a sample participant for each language (the same participants are used as those used in Figure 1). A significance map was generated for each participant by FreeSurfer(Dale et al., 1999); each map was smoothed using a Gaussian kernel of 4 mm full-width half-max and thresholded at the 70^th^ percentile of the positive contrast for each participant (this was done separately for each hemisphere). The surface overlays were rendered on the 80% inflated white-gray matter boundary of the fsaverage template using FreeView/FreeSurfer. Opaque red and yellow correspond to the 80^th^ and 99^th^ percentile of positive-contrast activation for each subject, respectively. Further, here and in Figure 1, small and/or idiosyncratic bits of activation (relatively common in individual-level language maps; e.g., Fedorenko et al., 2010; Mahowald & Fedorenko, 2016; Lipkin et al., in prep.-a) were removed. In particular, clusters were excluded if a) their surface area was below 100 mm^2, or b) they did not overlap (by >10%) with a mask created for a large number (n=804; Lipkin et al., in prep.-b) participants by overlaying the individual maps and excluding vertices that did not show language responses in at least 5% of the cohort. (We ensured that the idiosyncrasies were individual- and not language-specific: for each cluster removed, we checked that a similar cluster was not present for the second native speaker of that language.) These maps were used solely for visualization; all the statistical analyses were performed on the data analyzed in the volume.

**Supplementary Figure 2:**
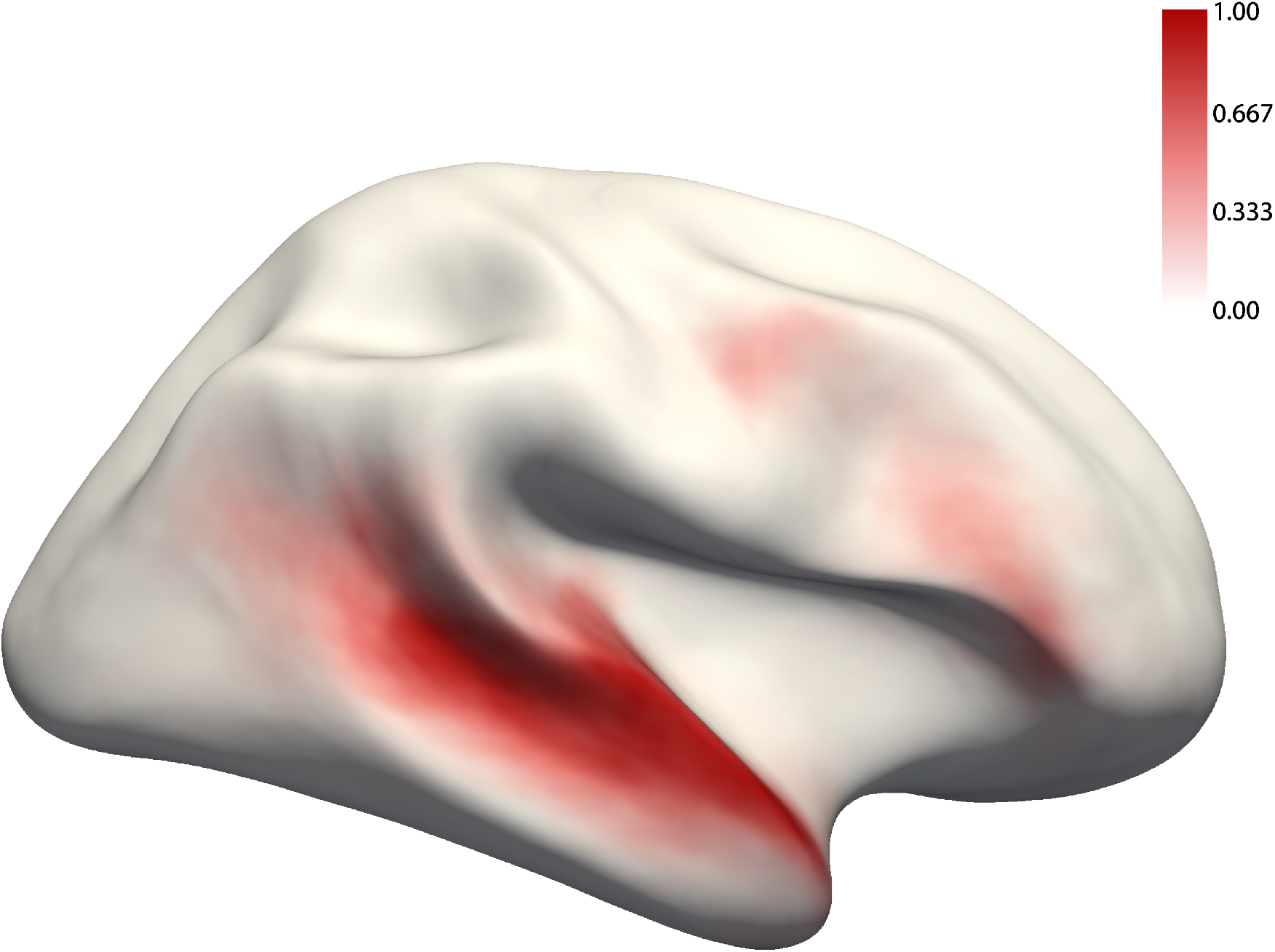
The probabilistic overlap map for the *Native-language>Degraded-language* contrast for the right hemisphere. This map was created by binarizing and overlaying the 86 participants’ individual maps (like those shown in Supp. Figure 1). The value in each vertex corresponds to the proportion of participants for whom that vertex belongs to the language network (see Supp. Figure 12 for a comparison between this probabilistic atlas vs. atlases based on native speakers of the same language).

**Supplementary Figure 3:**
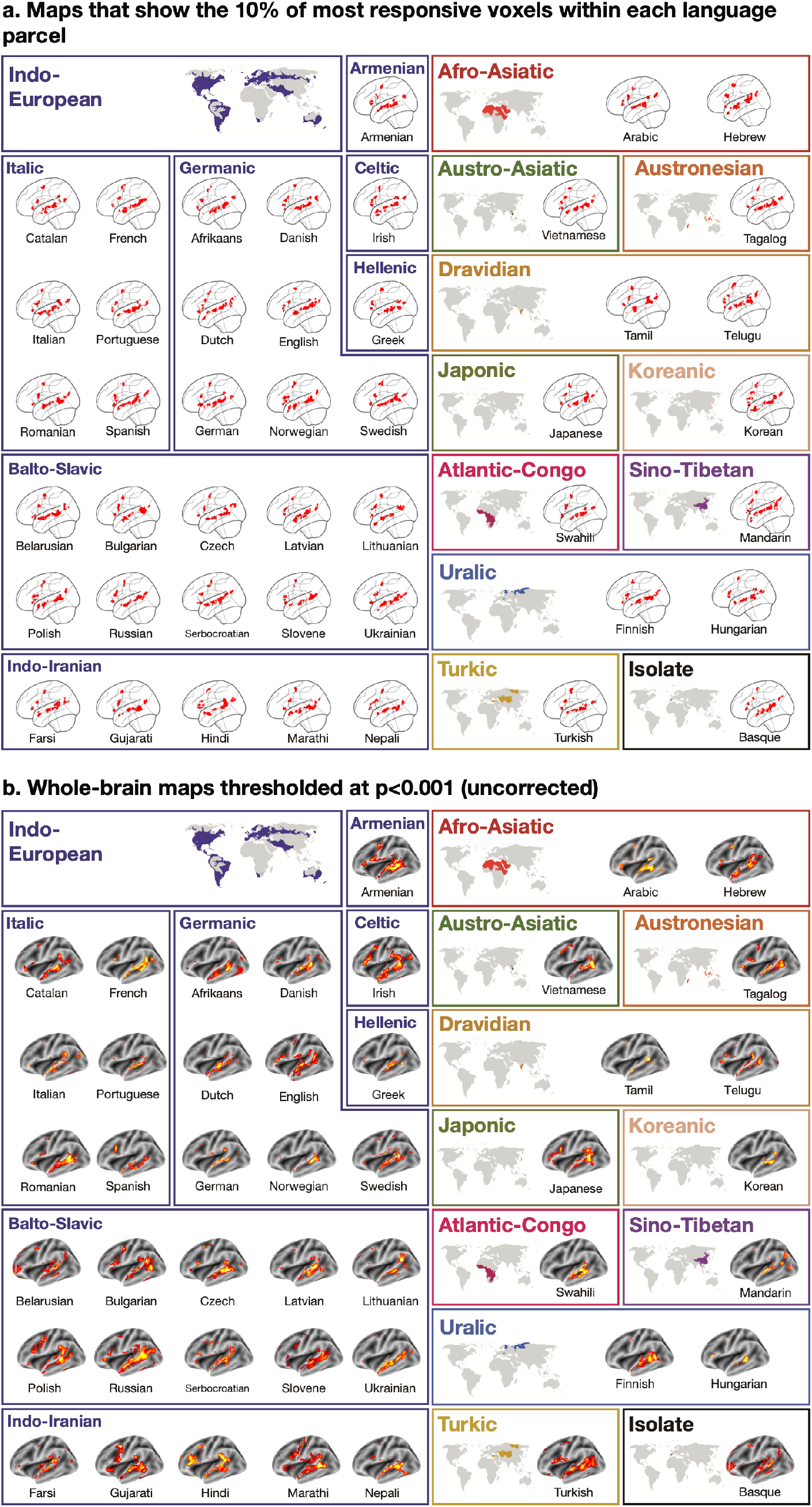
Volume-based activation maps for the *Native-language>Degraded-language* contrast in the left hemisphere of a sample participant for each language (the same participants are used as those used in Figure 1 and Supp. Figure 1). a) Binarized maps that were generated for each participant by selecting the top 10% most responsive (to this contrast) voxels within each language parcel. These sets of voxels correspond to the fROIs used in the analyses reported in Supp. Figure 4 (except for the estimation of the responses to the conditions of the Alice localizer, where a subset of the runs was used to ensure independence; the fROIs in those cases will be similar but not identical to those displayed). b) Whole-brain maps that are thresholded at the p<0.001 uncorrected level.

**Supplementary Figure 4:**
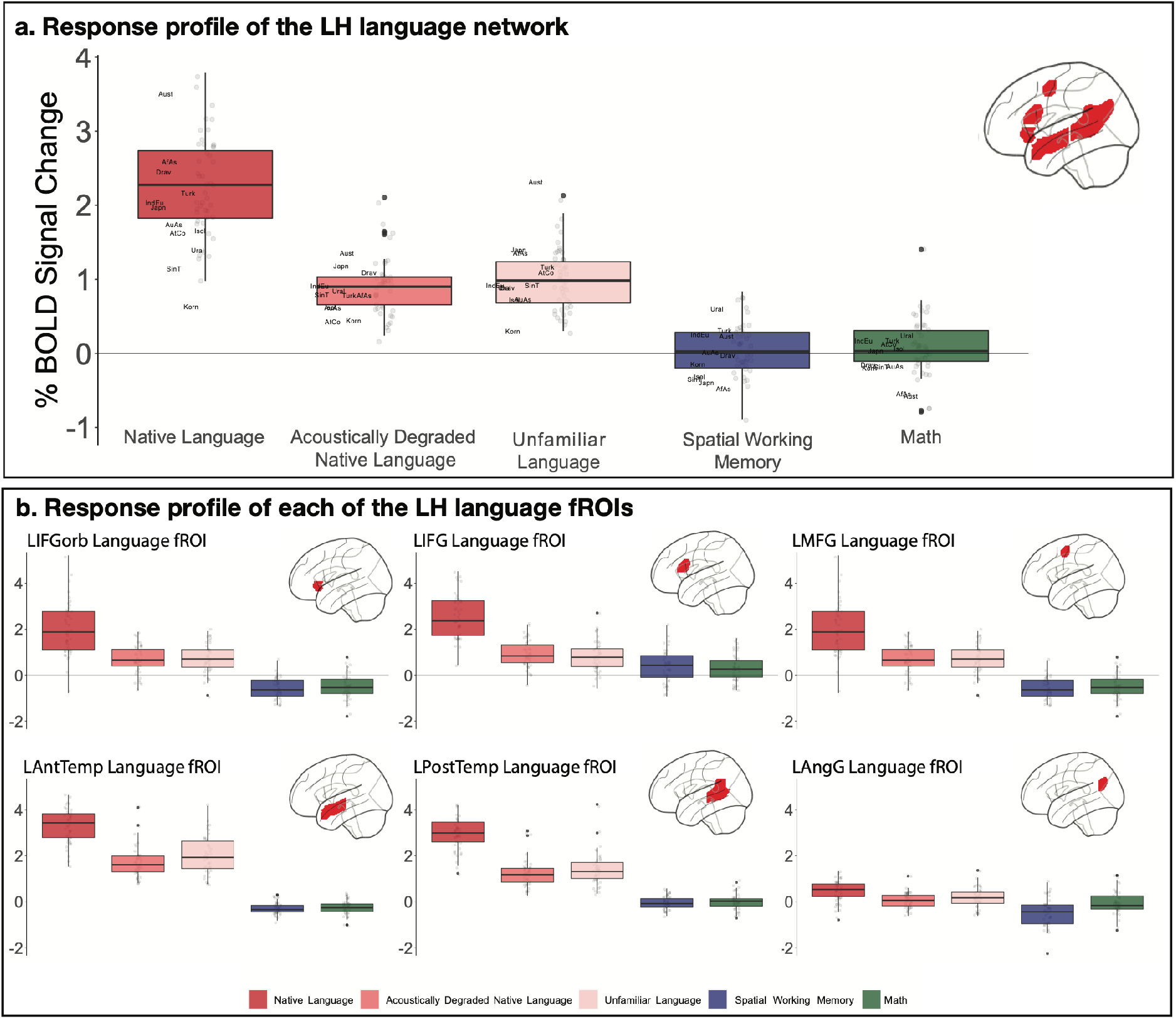
Percent BOLD signal change across (panel a) and within each of (panel b) the LH language functional ROIs (defined by the *Native-language>Degraded-language* contrast from the Alice localizer, cf. the *Sentences>Nonwords* contrast from the English localizer as in the main text and analyses; Figure 3a and Supp. Figure 7) for the three language conditions of the Alice localizer task (Native language, Acoustically degraded native language, and Unfamiliar language), the spatial working memory (WM) task and the math task. The dots correspond to languages (n=45), and the labels (panel a only) mark the averages for each language family. Across the six fROIs, the *Native-language* condition elicits a reliably greater response than both the *Degraded-language* condition (2.32 vs. 0.91 % BOLD signal change relative to the fixation baseline; t(44)=18.57, p<0.001) and the *Unfamiliar-language* condition (2.32 vs. 0.99; t(44)=18.02, p<0.001). Responses to the *Native-language* condition are also significantly higher than those to the spatial working memory task (2.32 vs. 0.06; t(44)=11.16, p<0.001) and the math task (2.32 vs. −0.02; t(40)=20.8, p<0.001). These results also hold for each fROI separately, correcting for the number of fROIs (*Native-language* > *Degraded-language*: ps<0.05; *Native-language* > *Unfamiliar-language*: ps<0.05; *Native-language* > *Spatial WM*: ps<0.05; and *Native-language* > *Math*: ps<0.05).

**Supplementary Figure 5:**
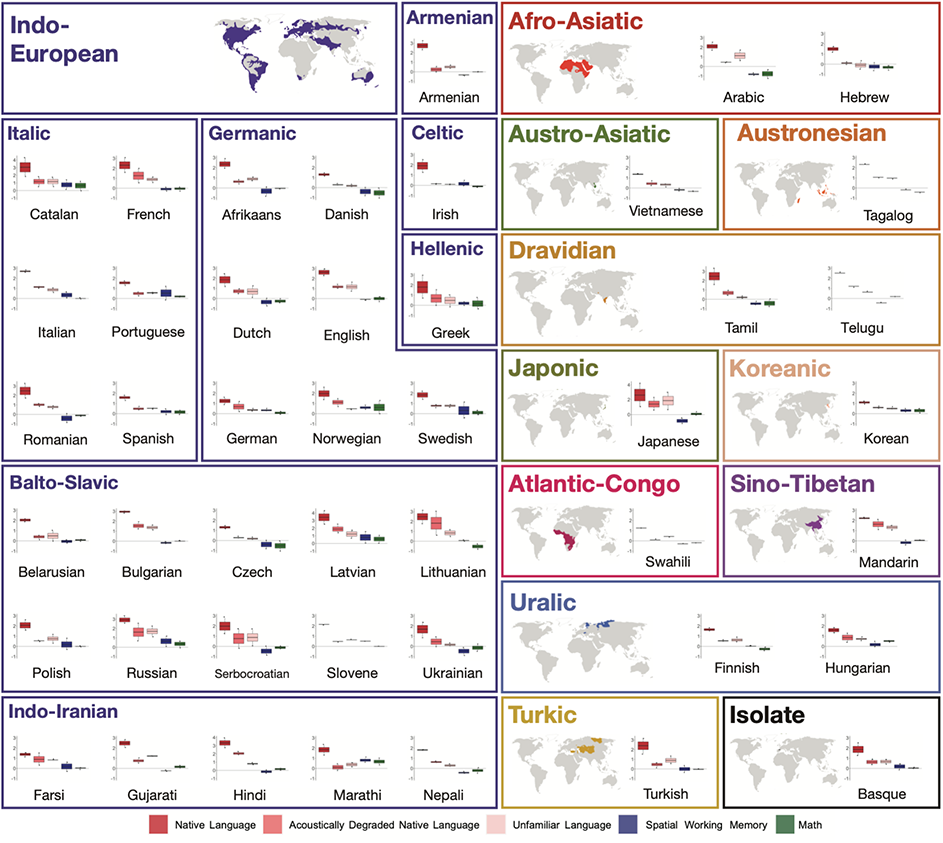
Percent BOLD signal change across the LH language functional ROIs (defined by the *Sentences>Nonwords* contrast) for the three language conditions of the Alice localizer task (Native language, Acoustically degraded native language, and Unfamiliar language), the spatial working memory (WM) task, and the math task shown for each language separately. The dots correspond to participants for each language. (Note that the scale of the y-axis differs across languages in order to allow for easier between-condition comparisons in each language.)

**Supplementary Figure 6:**
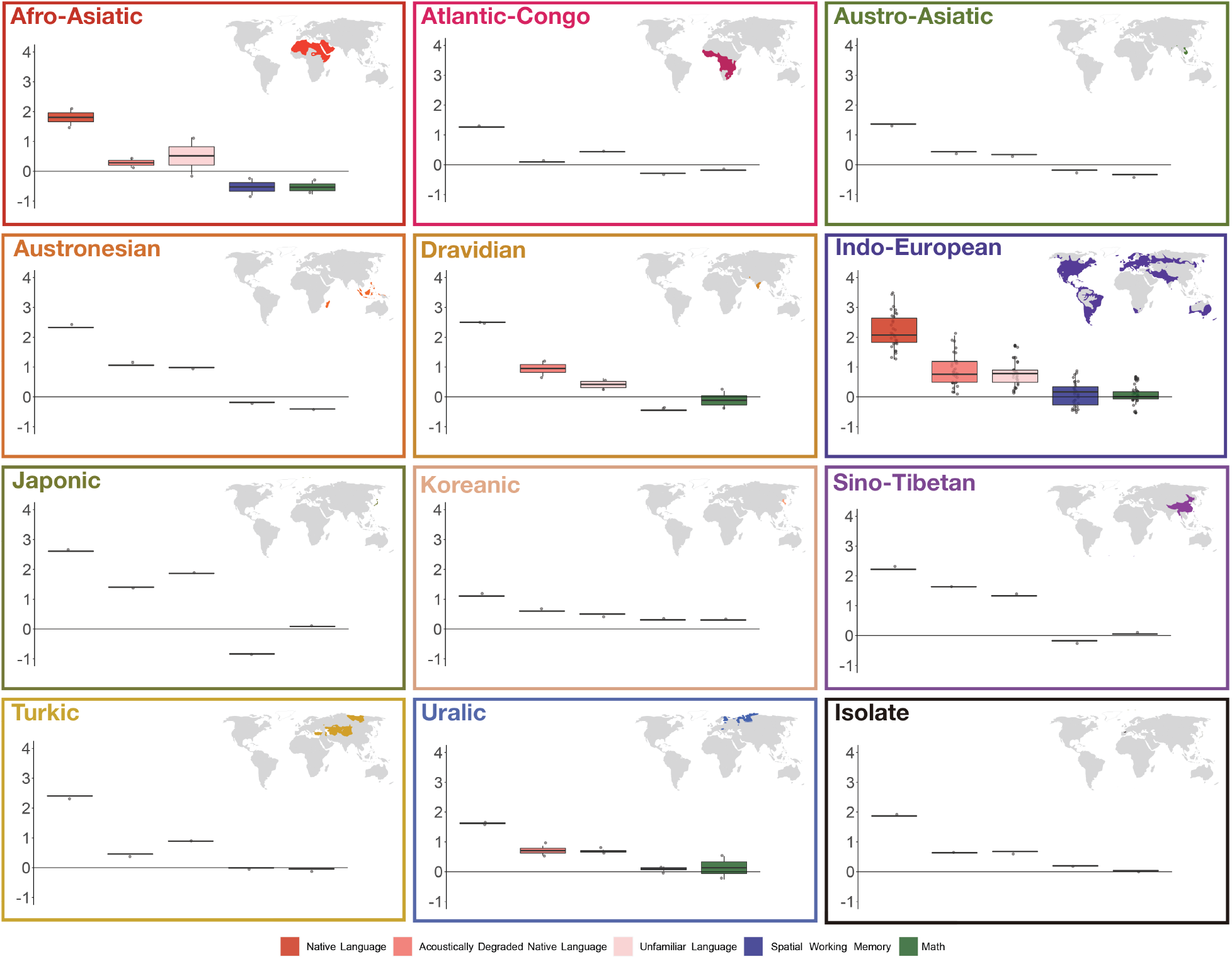
Percent BOLD signal change across the LH language functional ROIs (defined by the *Sentences>Nonwords* contrast) for the three language conditions of the Alice localizer task (Native language, Acoustically degraded native language, and Unfamiliar language), the spatial working memory (WM) task, and the math task shown for each language family separately. Across language families (n=12), the *Native-language* condition elicits a reliably greater response than both the *Degraded-language* condition (t(11)=9.92, p<0.001) and the *Unfamiliar-language* condition (t(11)=9.53, p<0.001). The *Native-language*>*Degraded-language* effect is stronger in the left hemisphere than the right hemisphere (t(11)=3.90, p=0.002), and more spatially extensive (t(11)=4.01, p<0.001). The regions of the LH language network exhibit strong correlations in their activity during story comprehension and rest, both reliably higher than zero (ts>4, ps<0.001) and phase-shuffled baselines (ts>10, ps<0.001). Further, the inter-region correlations in the LH language network are reliably stronger than those in the RH during both story comprehension (t(11)=4.06, p<0.01) and rest (t(11)=4.78, p<0.001). Responses to the *Native-language* condition are significantly higher than those to the spatial working memory task (t(11)=10.08, p<0.001) and the math task (t(11)=11.7, p<0.001). Furthermore, the language regions are dissociated in their intrinsic fluctuation patterns from the regions of the MD network: within-network correlations are reliably greater than between-network correlations both during story comprehension (ts>8, ps<0.001) and rest (ts>12, ps<0.001).

**Supplementary Figure 7:**
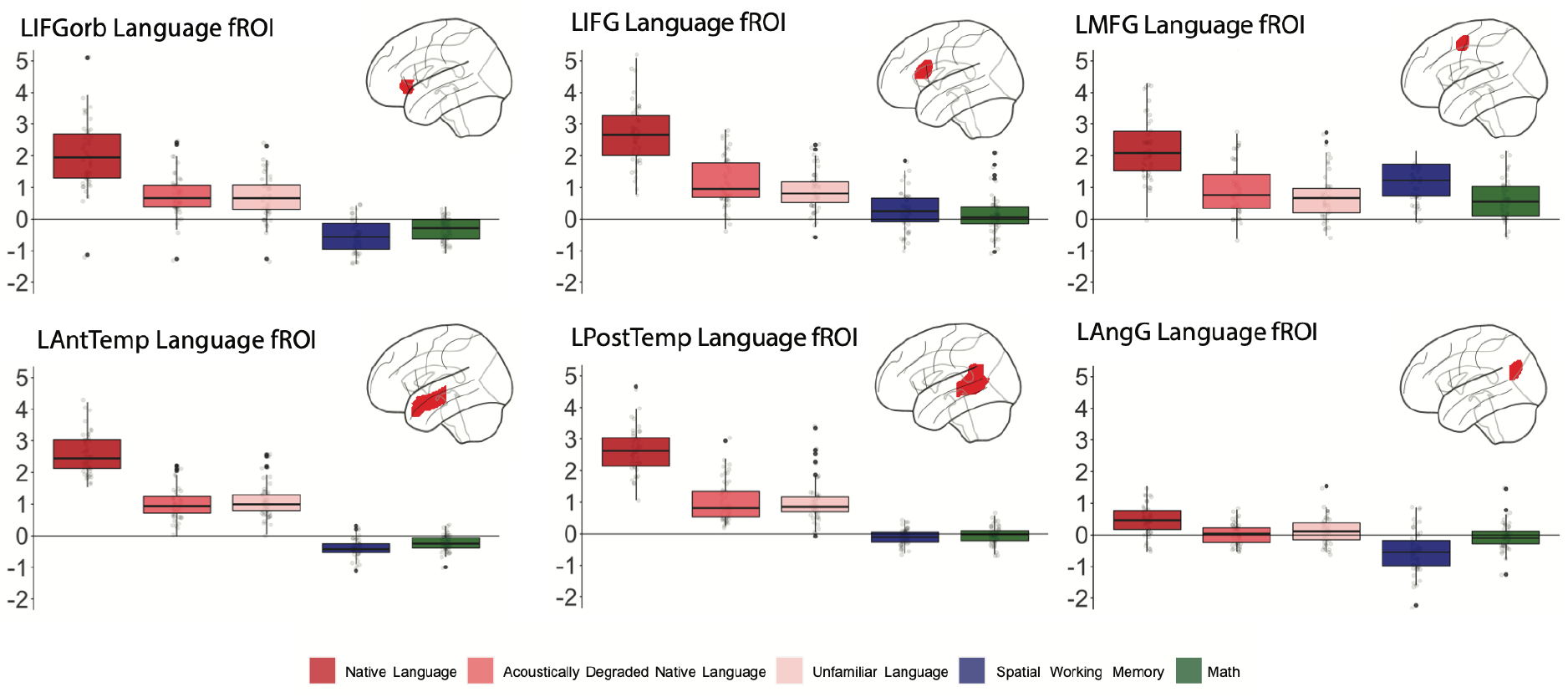
Percent BOLD signal change for each of the six LH language functional ROIs (defined by the *Sentences>Nonwords* contrast) for the three language conditions of the Alice localizer task (Native language, Acoustically degraded native language, and Unfamiliar language), the spatial working memory task, and the math task. The dots correspond to languages (n=45).

**Supplementary Figure 8:**
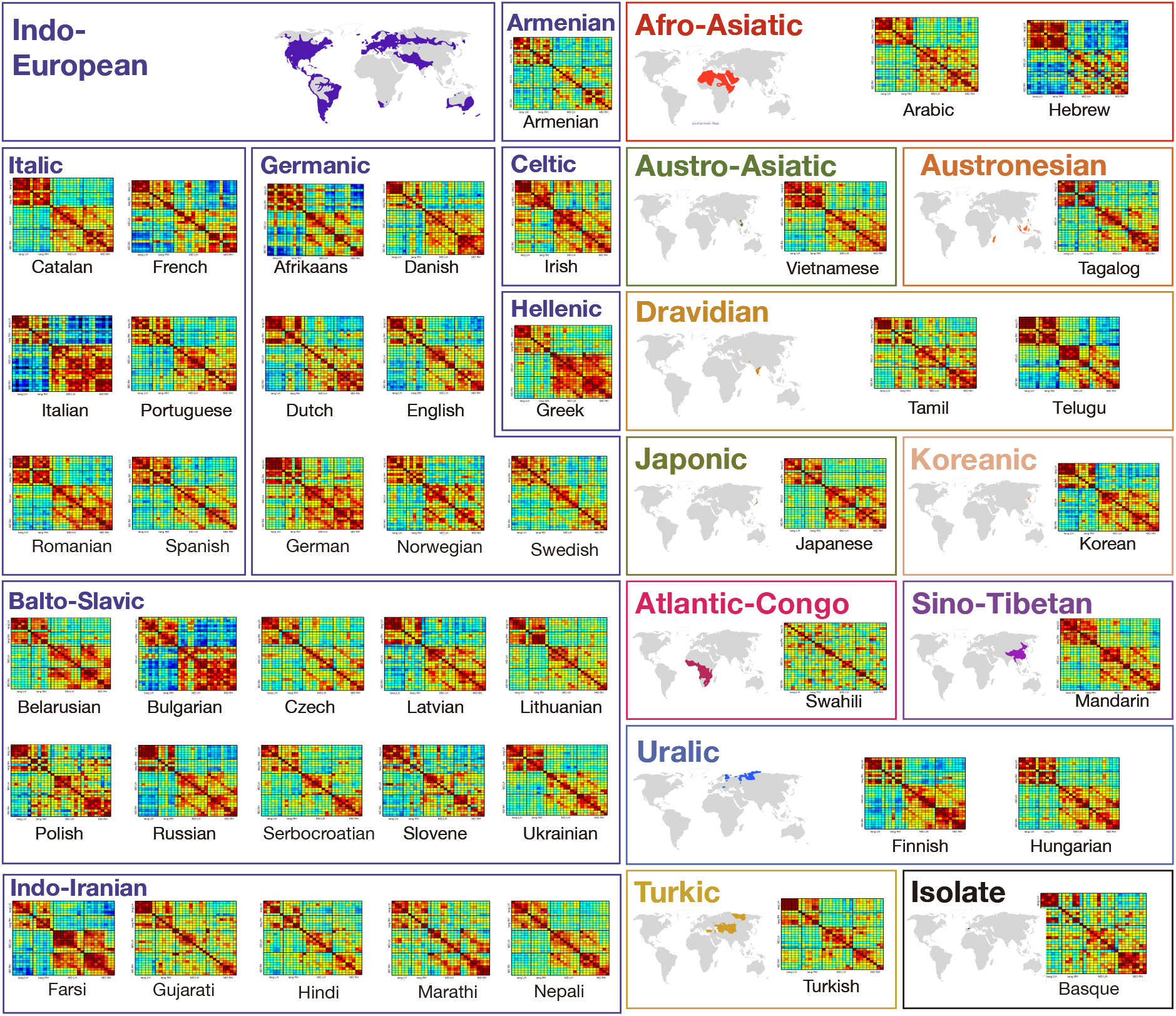
Inter-region functional correlations for the LH and RH of the language and the Multiple Demand (MD) networks during a naturalistic cognition paradigm (story comprehension in the participant’s native language) shown for each language separately.

**Supplementary Figure 9:**
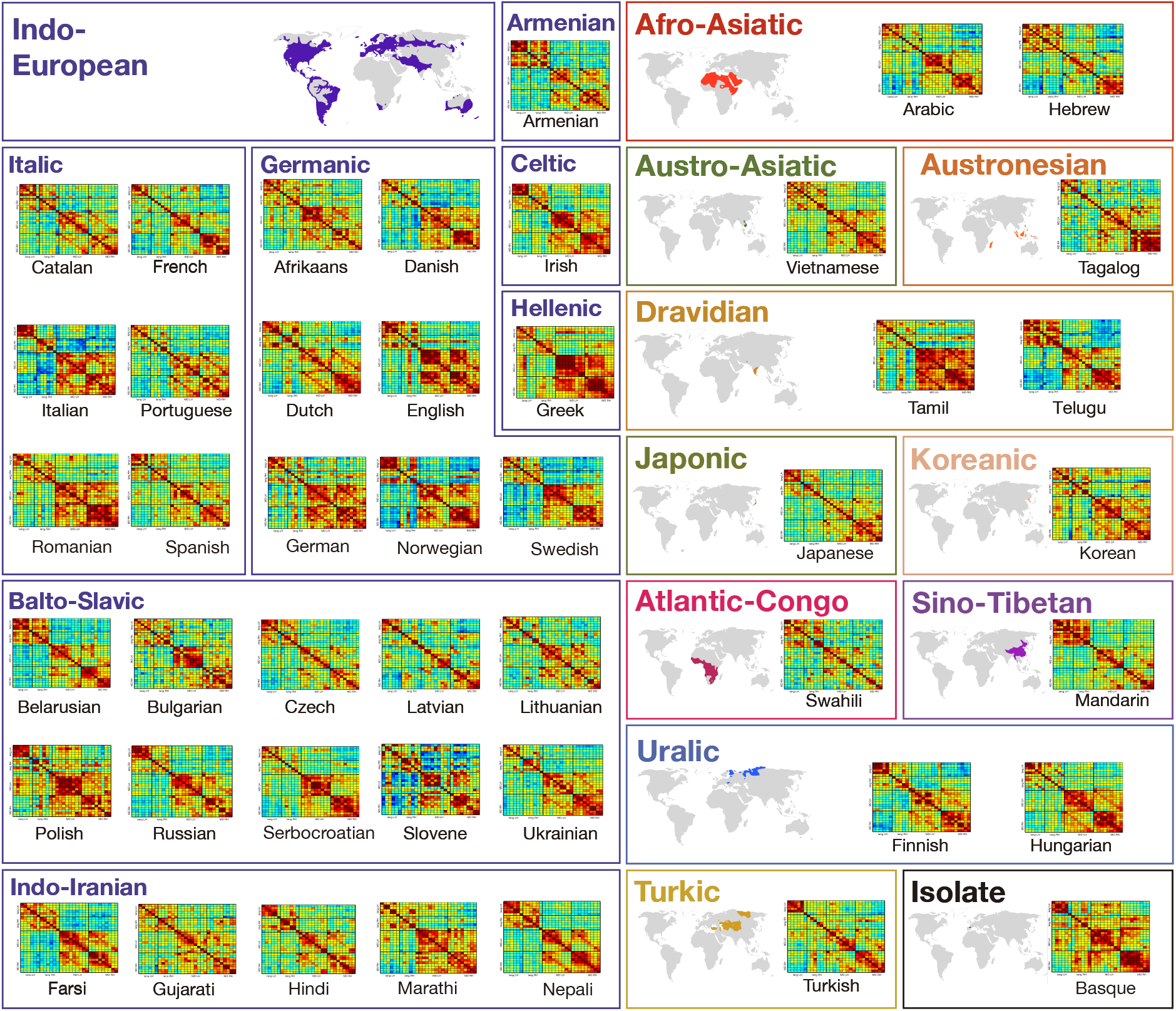
Inter-region functional correlations for the LH and RH of the language and the Multiple Demand (MD) networks during a naturalistic cognition paradigm (resting state) shown for each language separately.

**Supplementary Figure 10:**
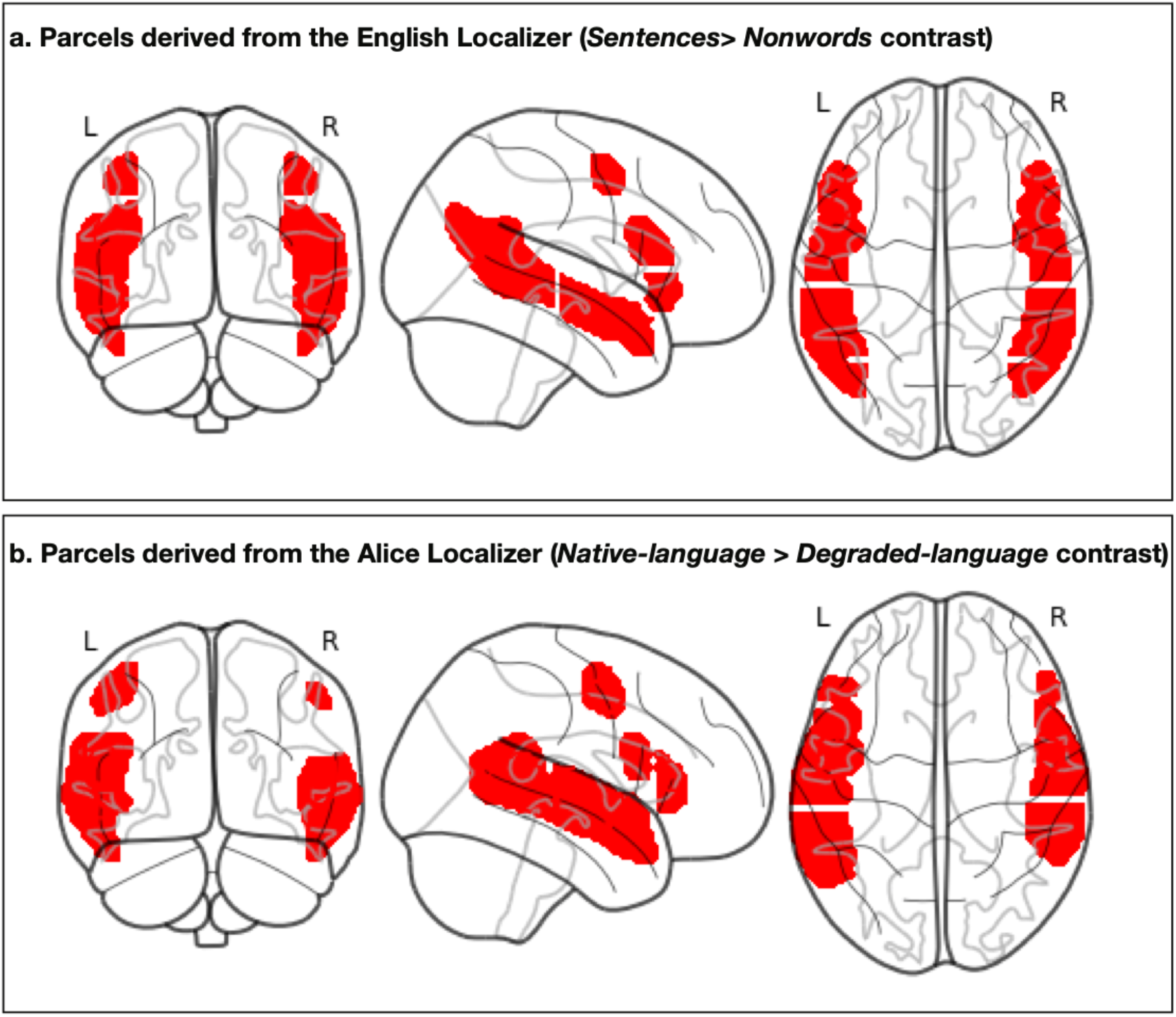
A visual comparison of the parcels that are used in the current study (derived via a Group-constrained Subject-Specific (GSS) approach(Fedorenko et al., 2010) from the probabilistic overlap map for the *Sentences>Nonwords* contrast in n=220 independent participants), and the parcels derived (also via GSS) from the probabilistic overlap map for the *Native-language>Degraded-language* contrast in the participants (n=86) in the current study. (Although the temporal-lobe parcels for the latter extend somewhat more superiorly, the *fROIs* selected based on contrasts between language and some perceptually-matched control condition—i.e., contrasts that target high-level language processing—are ∼identical for visual and auditory contrasts(Scott et al., 2017).)

**Supplementary Figure 11:**
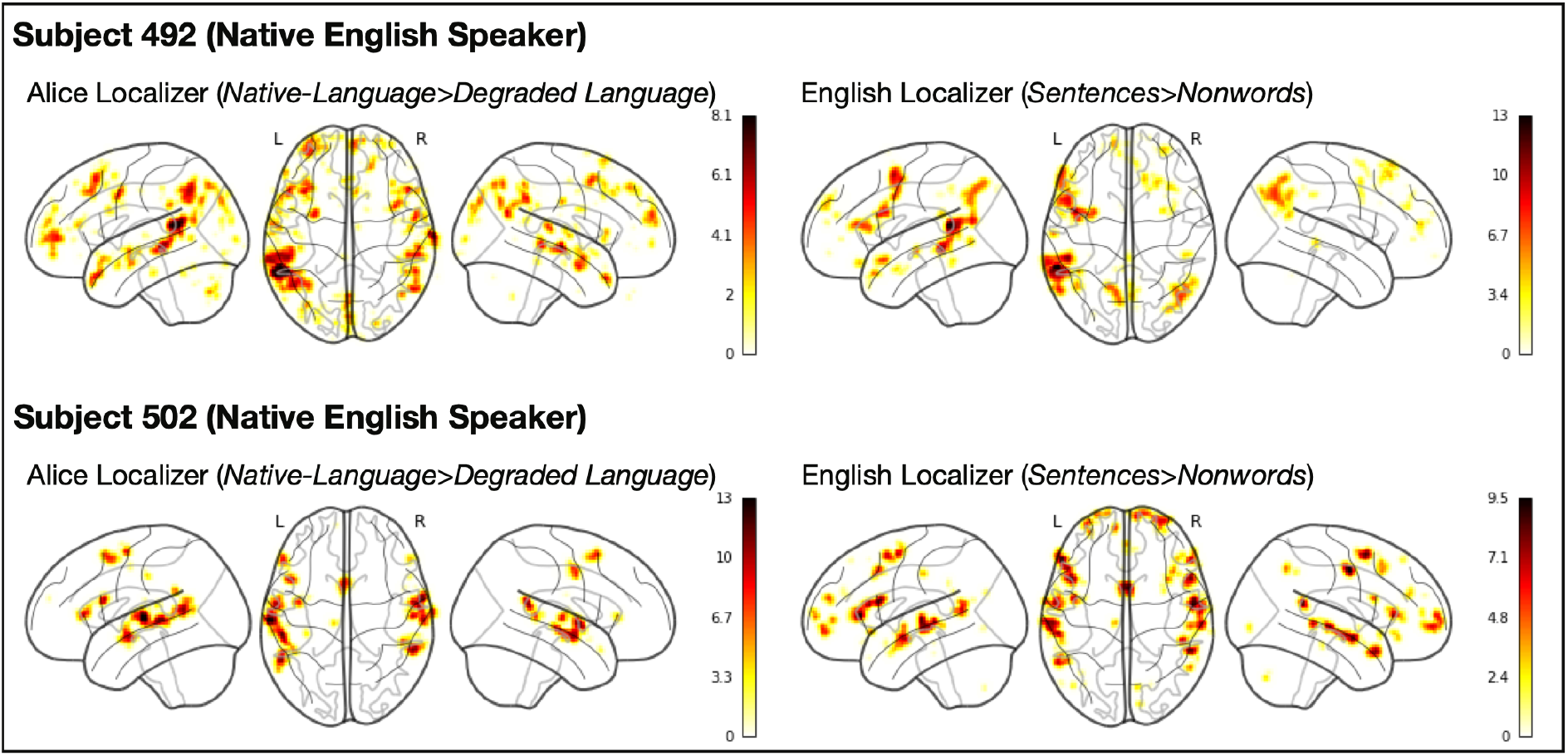
Comparison of the individual activation maps for the *Sentences>Nonwords* contrast and the *Native-language>Degraded-language* contrast in the two native-English-speaking participants. The two maps are voxel-wise (within the union of the language parcels) spatially correlated at r=0.77 and r=0.99 for participants 492 and 502, respectively (the correlations are Fisher-transformed). Across the full set of participants, the average Fisher-transformed spatial correlation between the maps for the *Sentences>Nonwords* contrast in English and the *Native-language>Degraded-language* contrast in the participant’s native language (again, constrained to the language parcels) is r=0.88 (SD=0.43) for the left hemisphere and 0.73 (SD=0.38) for the right hemisphere. (Note that using the union of the language parcels rather than the whole brain is conservative for computing these correlations; including all the voxels would inflate the correlations due to the large difference in activation levels between voxels that fall within the language parcels vs. outside their boundaries. Instead, we are zooming in on the activation landscape within the frontal, temporal, and parietal areas that house the language network and showing that these landscapes are spatially similar between the two contrasts in their fine-grained activation patterns.)

**Supplementary Figure 12:**
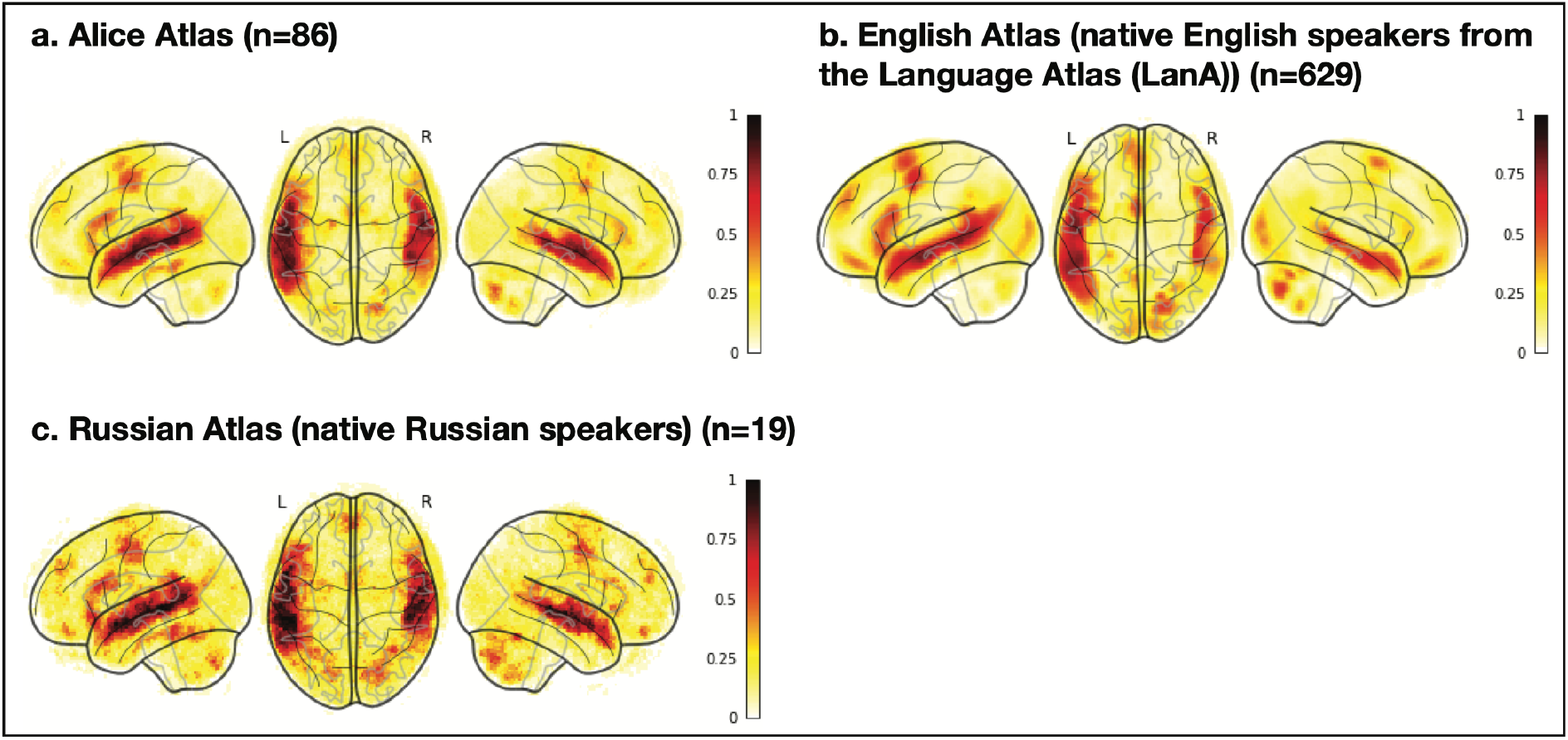
Comparison of three probabilistic overlap maps (atlases): a) the Alice atlas (n=86 native speakers of 45 languages) created from the *Native-language>Degraded-language* maps; b) the English atlas (n=629 native English speakers; this is a subset of the Fedorenko lab’s Language Atlas (LanA; Lipkin et al., in prep.-b)) created from the *Sentences>Nonwords* maps; and) the Russian Atlas (n=19 native Russian speakers) created from the *Native-language>Degraded-language* maps for the Russian version of the Alice localizer. All three atlases were created by selecting for each participant the top 10% of voxels (across the brain) based on the *t*-values for the relevant contrast in each participant, binarizing these maps, and then overlaying them in the common space. In each atlas, the value in each voxel corresponds to the proportion of participants (between 0 and 1) for whom that voxel belongs to the 10% of most language-responsive voxels. The probabilistic landscapes are similar across the atlases: within the union of the language parcels (see Supp. Figure 11 for an explanation of why this is more conservative than performing the comparison across the brain), the Alice atlas is voxel-wise spatially correlated with both the English atlas (r=0.83) and the Russian atlas (r=0.85). Furthermore, the range of positive overlap values is comparable between the Alice atlas (0.1-0.87; average within the language parcels=0.08, median=0.05) and each of the other atlases (the English atlas: 0.002-0.79; average within the language parcels=0.07, median=0.03; the Russian atlas: 0.05-0.84; average within the language parcels=0.13, median=0.11). The latter result suggests that the inter-individual variability in the topographies of activation landscapes elicited in 86 participants of 45 diverse languages is comparable to the inter-individual variability observed among native speakers of the same language.

**Supplementary Figure 13:**
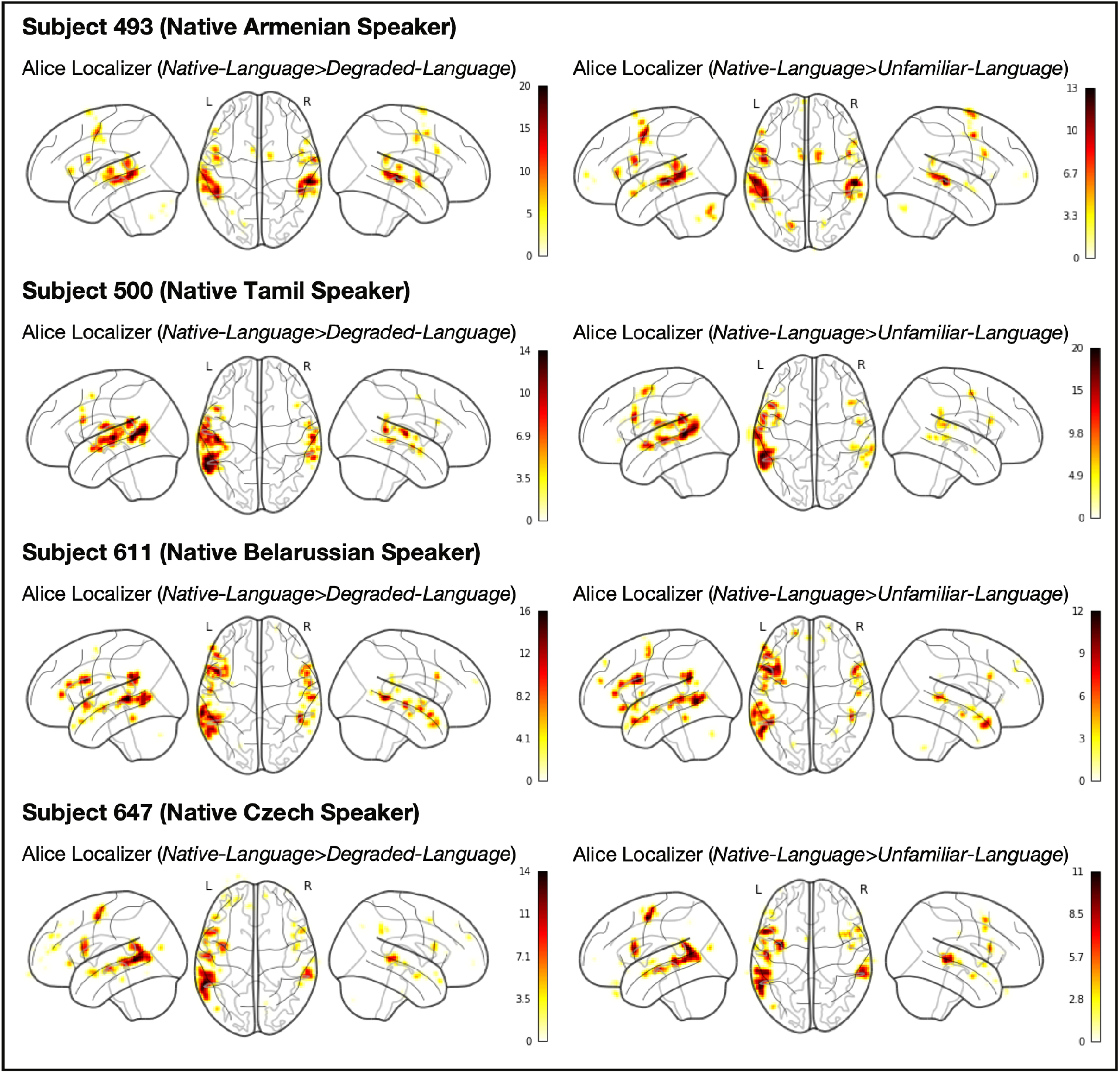
Comparison of the individual activation maps for the *Native-language*>*Degraded-language* contrast and the *Native-language*>*Unfamiliar-language* contrast in four sample participants. The activation landscapes are broadly similar: across the full set of 86 participants, the average Fisher-transformed voxel-wise spatial correlation within the union of the language parcels between the maps for the two contrasts is r=0.66 (SD=0.40). (Note that this correlation is lower than the correlation between the *Native-language*>*Degraded-language* contrast and the *Sentences>Nonwords* contrast in English (see Supp. Figure 11). This difference may be due to the greater variability in the participants’ responses to an unfamiliar language.) Furthermore, across the language fROIs, the magnitudes of the *Native-language*>*Degraded-language* and the *Native-language*>*Unfamiliar-language* effects are similar (mean = 1.02, SD(across languages)=0.41 vs. mean=1.07, SD=0.37, respectively; t(44)=1.15, p=0.26).

**Supplementary Figure 14:**
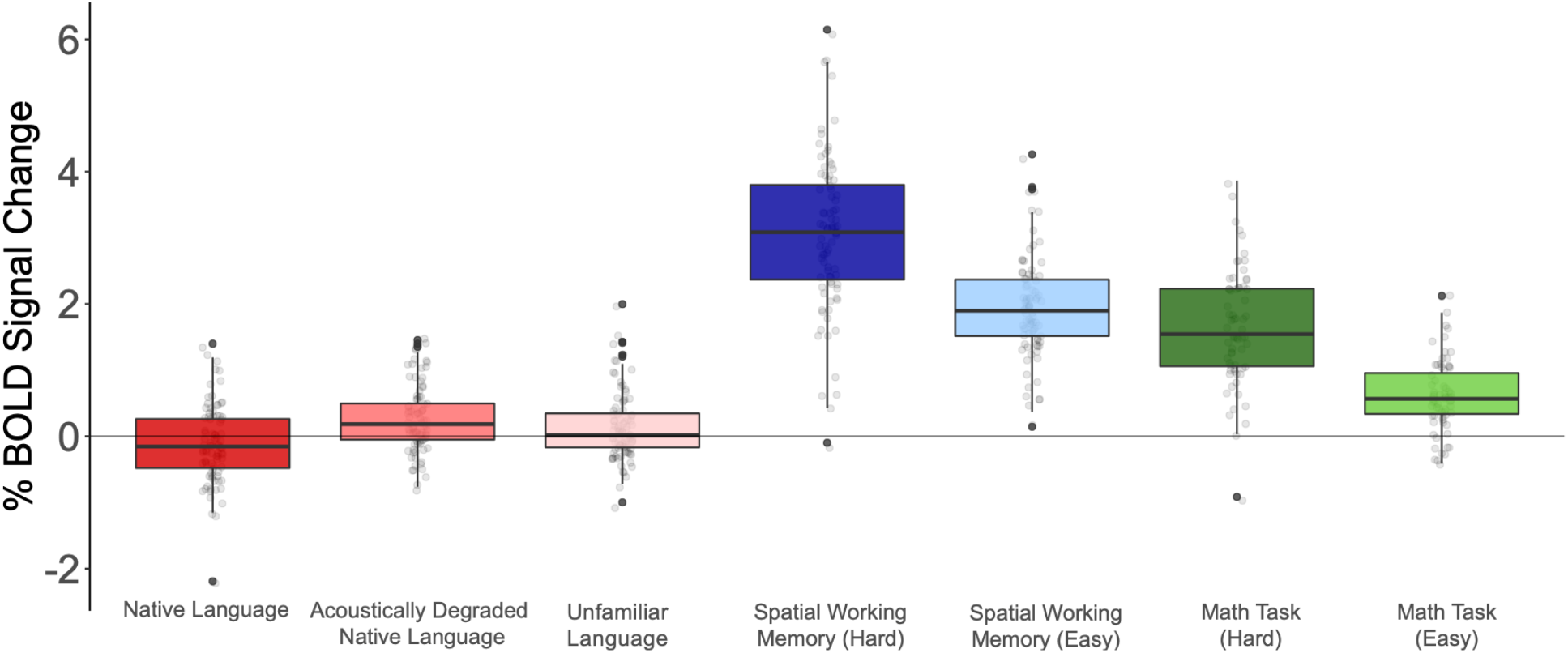
Percent BOLD signal change across the domain-general Multiple Demand (MD) network (Duncan, 2010, 2013) functional ROIs for the three language conditions of the Alice localizer task (Native language, Acoustically degraded native language, and Unfamiliar language), the hard and easy conditions of the spatial working memory (WM) task, and the hard and easy conditions of the math task. As in the main analyses (Figure 3c), the individual MD fROIs were defined by the *Hard>Easy* contrast in the spatial WM task (see Fedorenko et al., 2013 for evidence that other *Hard>Easy* contrasts activate similar areas). As expected given past work (e.g., Fedorenko et al., 2013), the MD fROIs show strong responses to both the spatial WM task and the math task, with stronger responses to the harder condition in each (3.05 vs. 1.93 for the spatial WM task, t(44)=23.1, p<0.001; and 1.68 vs. 0.62 for the math task, t(40)=8.87, p<0.001). These robust responses in the MD network suggest that the lack of responses to the spatial WM and math tasks in the language areas can be meaningfully interpreted. Furthermore, in line with past work (e.g., Davis et al., 2003; Hervais-Adelman et al., 2012; Erb et al., 2013), MD fROIs show a stronger response to the acoustically degraded condition than the native language condition (0.26 vs. −0.10, t(44)=4.92, p<0.01), and to the unfamiliar language condition than the native language condition (0.15 vs. −0.10, t(44)=4.96, p<0.01).

**Supplementary Figure 15:**
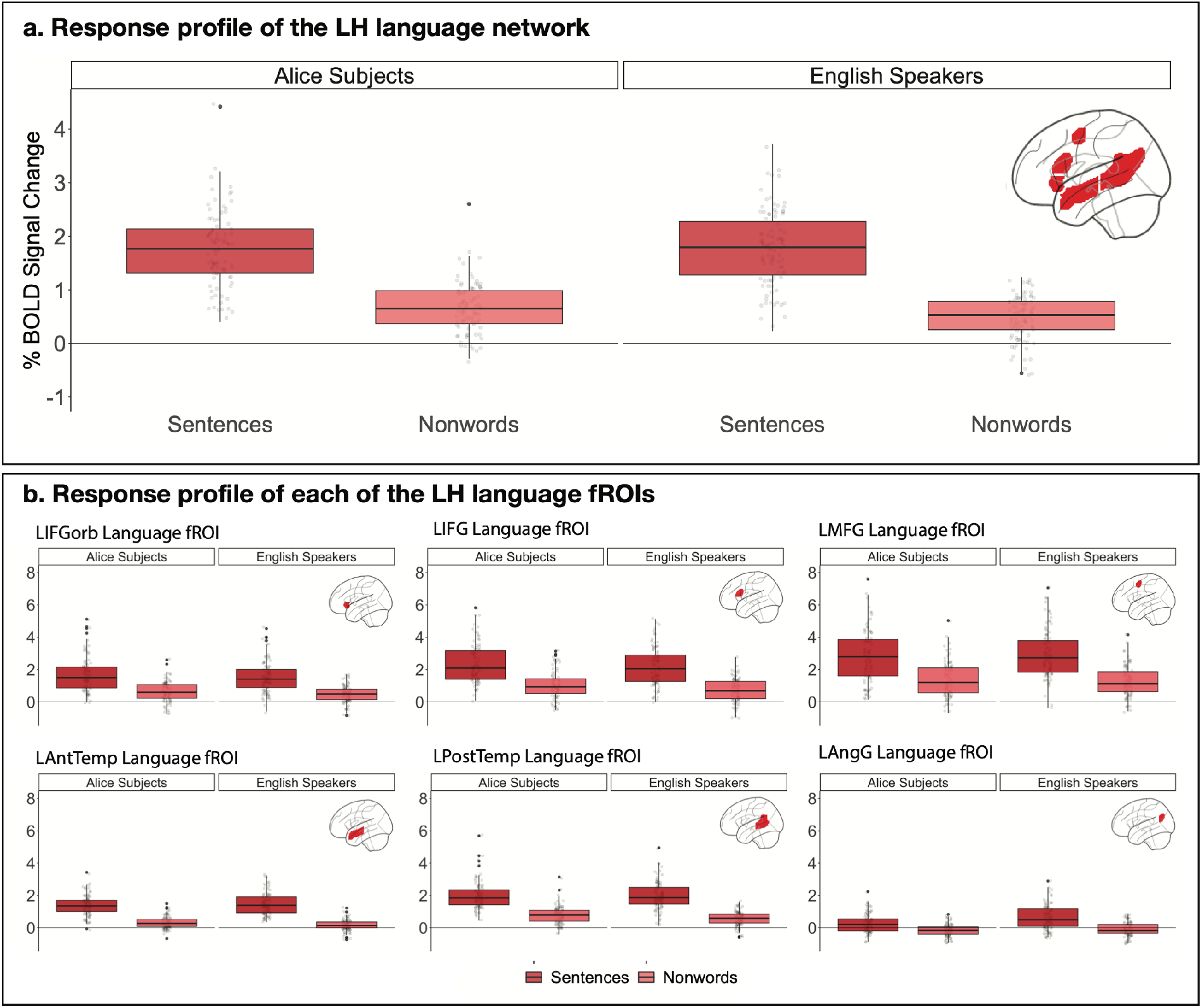
Percent BOLD signal change across (panel a) and within each of (panel b) the LH language functional ROIs (defined by the *Sentences>Nonwords* contrast; responses were estimated using across-runs cross-validation (Nieto-Castañón & Fedorenko, 2012), to ensure independence) for the Sentences and Nonwords conditions. The Alice subjects are the 86 participants from the current study (84 of whom are non-native but proficient speakers of English; we included the two native English speakers here for ease of comparing these results to the results in the rest of the paper where we report the results for the full set of 86 participants); the English speakers are a set of n=74 native English speakers (all learned English before the age of 5). The dots correspond to individual participants. Across the six LH fROIs, the *Sentences* condition elicits a reliably greater response than the *Nonwords* condition in both the Alice subjects (1.23 vs. 0.49 % BOLD signal change relative to the fixation baseline; t(85)=20.38, p<0.001) and the native English speakers (1.22 vs. 0.37; t(73)=18.8, p<0.001). The magnitude of response for the sentences condition is almost identical between the two populations (1.23 vs. 1.22, t<1); the magnitude of response for the nonwords condition is a little higher in the Alice subjects (0.49 vs. 0.37; t(157.36)=2.1, p=0.03). Because this difference was not predicted, we do not attempt to interpret it. Critically, this supplementary analysis shows that the response during the processing of English is similar between our Alice subjects and a set of native English speakers, and the *Sentences>Nonwords* contrast is similarly robust, suggesting that the use of this contrast as a language localizer is justified (as is also clear from Supp. Figure 4, which shows that similar responses obtain when the fROIs are defined by one’s native language localizer).

**Supplementary Figure 16:**
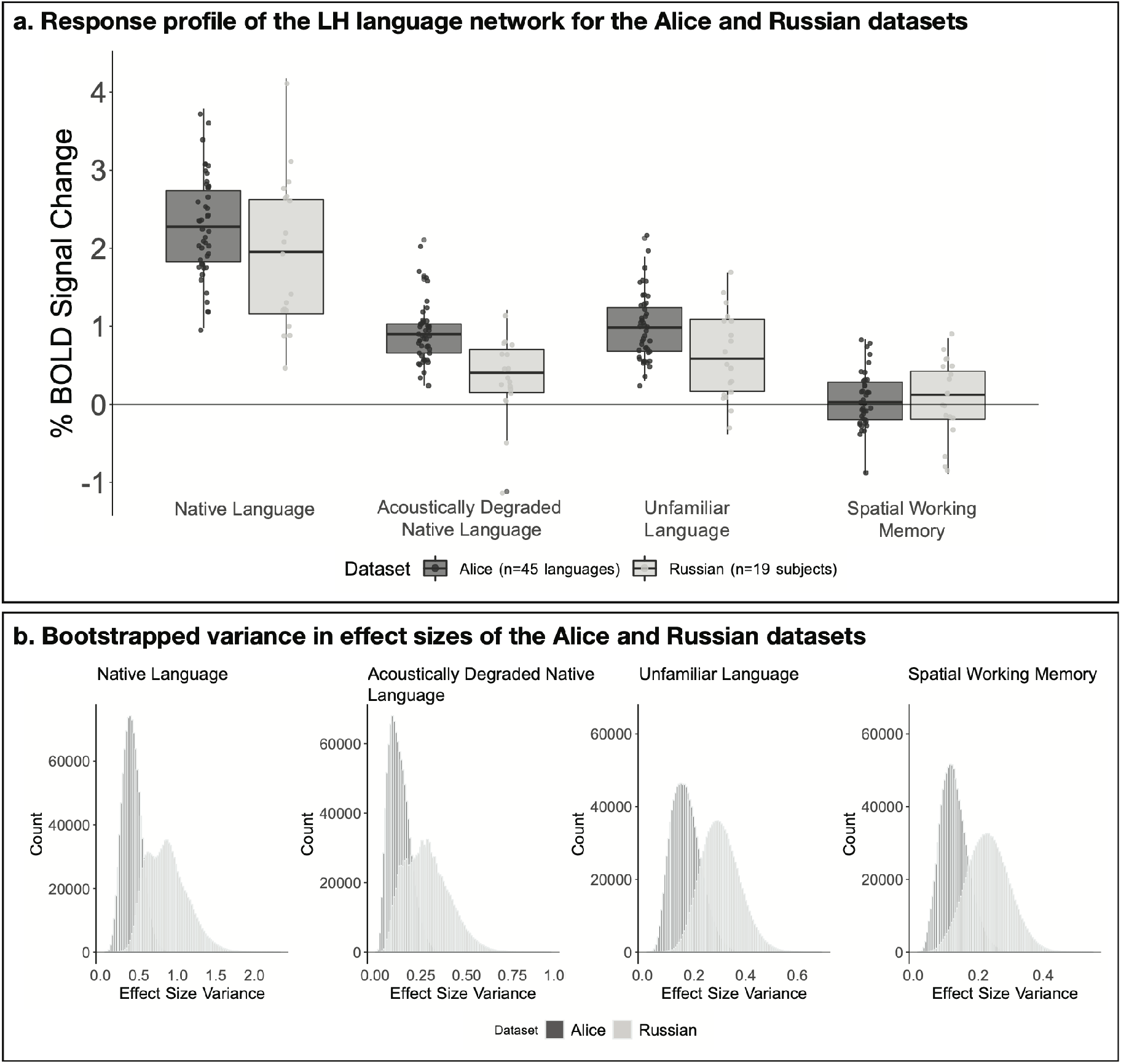
A comparison of inter-individual variability in effect sizes for one’s native language and the control conditions for speakers of diverse languages vs. for speakers of the same language (Russian). As can be seen in Figure 3a in the main text, we observed substantial variability across languages in the strength of neural response during language processing (and the control conditions). In order to compare the level of cross-linguistic variability to inter-individual variability for speakers of the same language (e.g., Mahowald & Fedorenko, 2016; Mineroff, Blank et al., 2018), we leveraged an existing dataset of 19 native speakers of Russian (see also Supp. Figure 12), who completed the Alice localizer (and the spatial working memory task included here for completeness; as in the main paper, we are averaging the responses across the hard and easy conditions). a) Percent BOLD signal change across the LH language functional ROIs (defined by the *Native-language>Degraded-language* contrast) for the three language conditions of the Alice localizer task (Native language, Acoustically degraded native language, and Unfamiliar language), and the spatial working memory (WM) task. Left bars (within each of the four conditions): the current dataset (n=45 languages (1-2 participants per language); dots=languages); right bars: a dataset of n=19 native Russian speakers (unfamiliar language = Tamil) (dots=individual participants). Visual inspection of the distributions of the individual data points suggests that cross-linguistic and inter-individual variability are comparable. b) Bootstrapped variance in effect sizes of the Alice dataset (n=86 participants) and the Russian dataset (n=19 participants) for each of the conditions in the Alice localizer task (Native language, Acoustically degraded native language, and Unfamiliar language). To perform this analysis, we bootstrapped (n=1,000,000) the effect sizes for each of the three conditions for the 86 participants in the Alice dataset (sampling 19 participants at a time) and for the 19 participants in the Russian dataset. If cross-linguistic variability is greater than the variability that exists among individual speakers of the same language, we should see higher variance in the Alice dataset compared to the Russian dataset. Instead, as can be seen in panel b, the variance in the Alice dataset is actually lower than that in the Russian dataset for the Native language condition (p=0.04), the Acoustically degraded condition (p=0.12), the Unfamiliar language condition (p=0.07), and the spatial working memory task (p=0.09). (The reason for numerically higher variability in the Russian dataset may have to do with a wider age range in that group.) As a result, the variability that we observe in the main Figure 3a likely reflects inter-individual rather than cross-linguistic variability. As discussed in the main text, however, future work may discover cross-linguistic differences (when a deep sampling approach is used, with large numbers of speakers tested for each language/language family)—in the measures examined here or some other ones—that would exceed inter-individual variability.

**Supplementary Figure 17:**
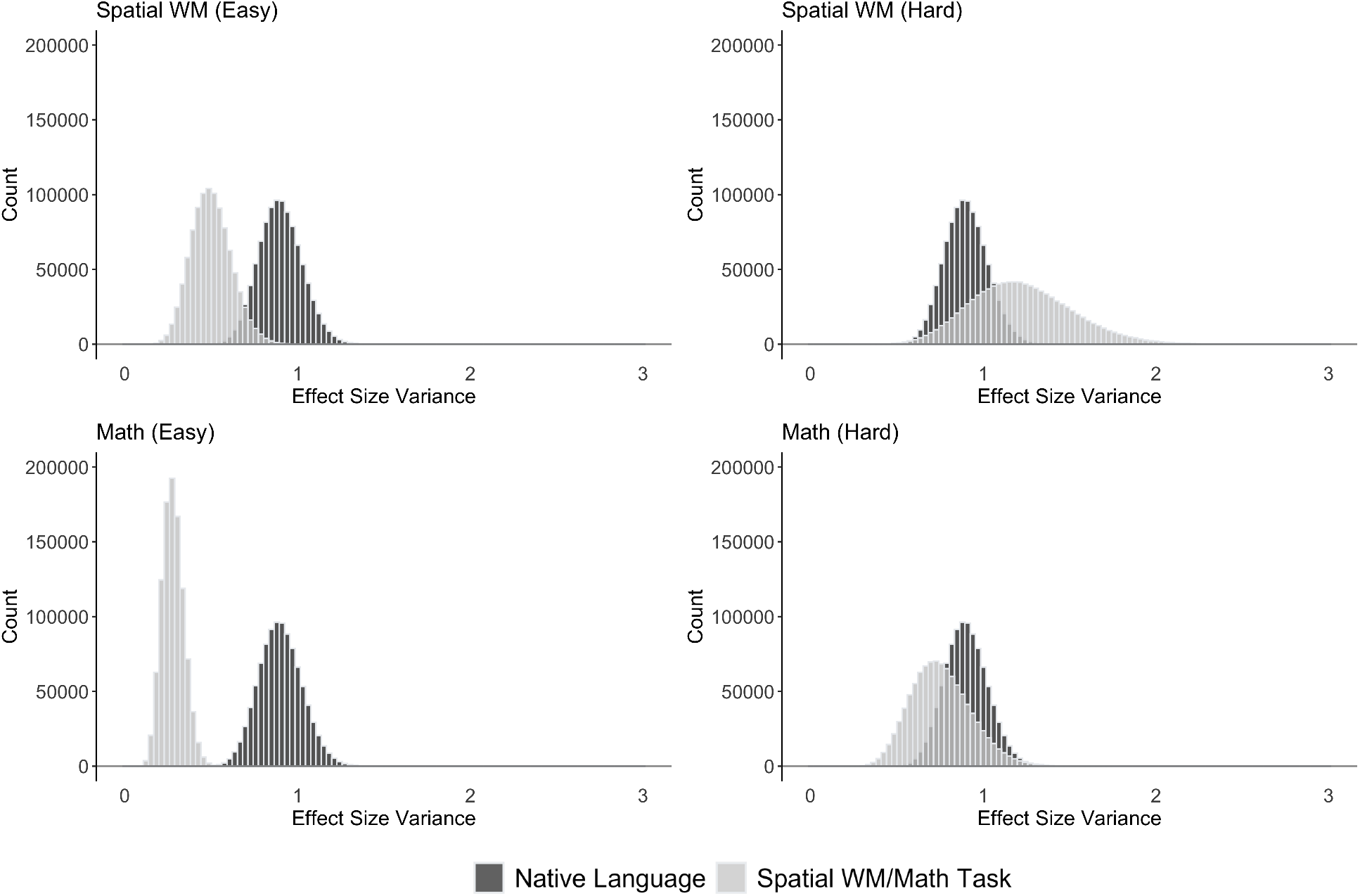
A comparison of inter-individual variability in effect sizes during language processing and during non-linguistic cognitive tasks for the Alice dataset (n=86 participants). Bootstrapped variance in effect sizes for the Native language condition in the Alice localizer task (dark grey; same distribution across the four panels) and the non-linguistic control task (light grey; top: spatial WM task, bottom: math task; left: easy condition, right: hard condition). To perform this analysis, we bootstrapped (n=1,000,000) the effect sizes in the LH language network for the Native language condition in the Alice localizer task, and in the bilateral MD network for each of the four non-linguistic conditions (which were identical across participants, in contrast to the Alice localizer task, which differed depending on the participant’s native language). If cross-linguistic variability is greater than the variability that exists in the strength of neural responses during non-linguistic tasks, we should see higher variance in response to the Native language condition compared to the responses to the different non-linguistic tasks, assuming the effect sizes are comparable (given that variance scales with effect sizes, we would generally expect to see higher variance for larger effects). This analysis revealed that the variance for the Native language condition was similar to the variance in the hard conditions—which elicit a strong response in the MD network—in both the spatial WM (p=0.75) and math (p=0.85) tasks, but was higher than the variance in the easy conditions—which elicit a relatively lower response in the MD network—in both the spatial WM (p=0.01) and math (p<0.01) tasks. The fact that the variance during native language processing is similar to the variance in the hard conditions suggests that the former is likely due to inter-individual variability rather than cross-linguistic variability.

**Supplementary Figure 18:**
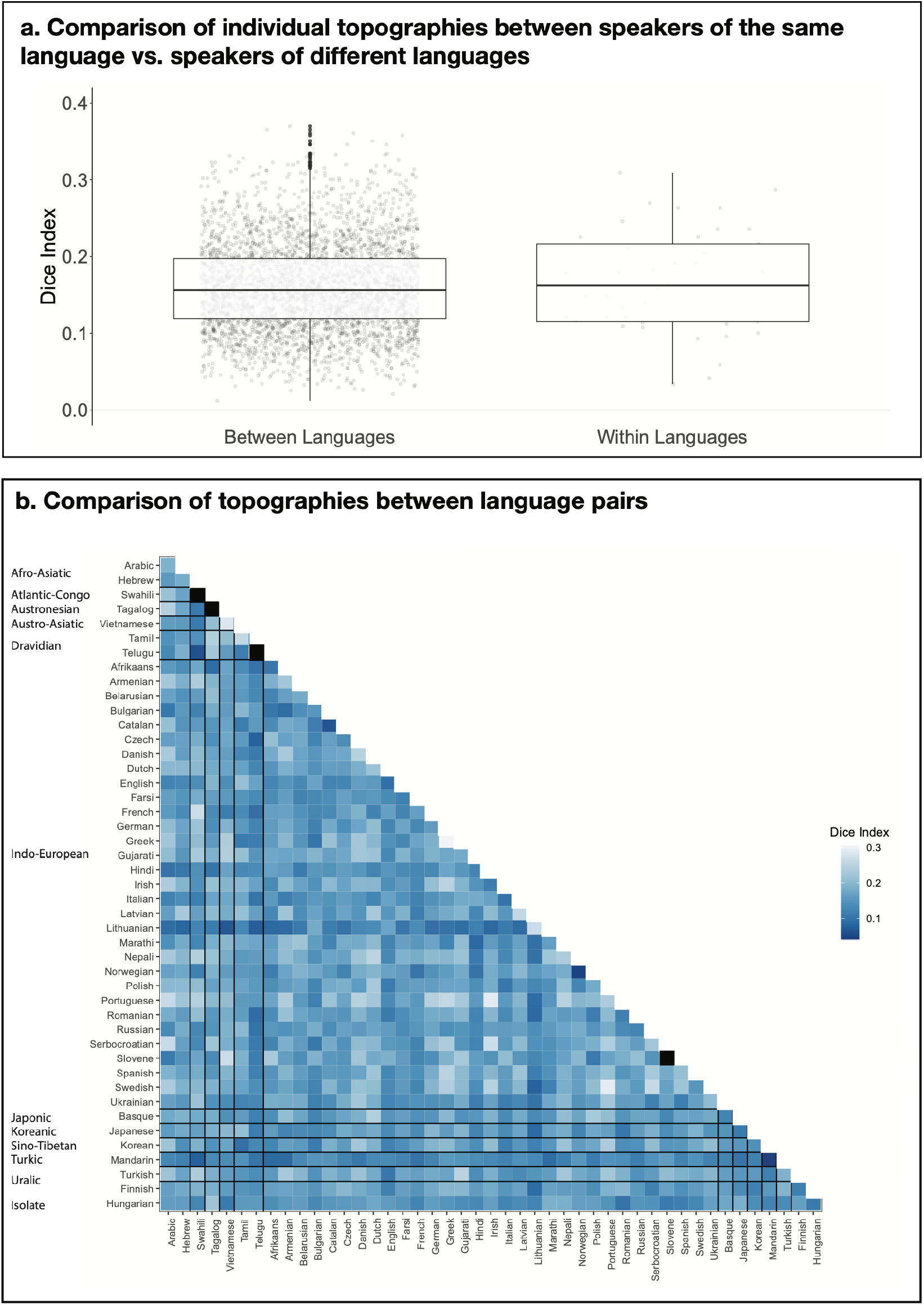
A comparison of individual LH topographies between speakers of the same language vs. between speakers of different languages. The goal of this analysis was to test whether inter-language / inter-language-family similarities might be reflected in the similarity structure of the activation patterns. To perform this analysis, we computed a Dice coefficient (Rombouts et al., 1997) for each pair of individual activation maps for the *Intact-language>Degraded-language* contrast (a total of n=3,655 pairs across the 86 participants). To do so, we used the binarized maps like those shown in Supp. Figure 3a, where in each LH language parcel top 10% of most responsive voxels were selected. Then, for each pair of images, we divided the number of overlapping voxels multiplied by 2 by the sum of the voxels across the two images (this value was always the same and equaling 1,358 given that each map had the same number of selected voxels). The resulting values can vary from 0 (no overlapping voxels) to 1 (all voxels overlap). a) A comparison of Dice coefficients for pairs of maps between languages (left) vs. within languages (right; this could be done for 41/45 languages for which two speakers were tested). If the activation landscapes are more similar within than between languages, then the Dice coefficients for the within-language comparisons should be higher. Instead, no reliable difference was observed by an independent-samples t-test (average within-language: 0.17 (SD=0.07), average between-language: 0.16 (SD=0.06); t(40.7)=-0.52, p= 0.61; see also Supp. Figure 12 for evidence that the range of overlap values in probabilistic atlases created from speakers of diverse languages vs. speakers of the same language are comparable). b) Dice coefficient values for all pairs of within- and between-language comparisons (the squares in black on the diagonal correspond to languages with only one speaker tested). As can be seen in the figure and in line with the results in panel a, no structure is discernible that would suggest greater within-language / within-language-family topographic similarity. Similar to the results from the within- vs. between-language comparison in a, the within-language-family vs. between-language-family comparison did not reveal a difference (t(19.8)=0.71, p=0.49). In summary, in the current dataset (collected with the shallow sampling approach, i.e., a small number of speakers from a larger number of languages), no clear similarity structure is apparent that would suggest more similar topographies among speakers of the same language, or among speakers of languages that belong to the same language family.

### Supplementary Tables

**Supplementary Table 1:** A partial selective review of past fMRI studies on the languages included in the current investigation. For each language, SMM performed searches (on Google, GoogleScholar, PubMed, etc.) for “fMRI [language]” (e.g., fMRI Ukranian) and extracted the relevant citations where available. All papers dealing with speech (perception and articulation), reading, and language (comprehension and production) were considered (i.e., we did not restrict our search to only papers that focus on high-level linguistic processing). Further, we included papers from the clinical literature (that simply used the language in question to facilitate pre-surgical planning rather than asking scientific questions about the particular language or language processing mechanisms in general) and papers where the language in question was used as a control condition. We classified languages into three groups: well studied languages (with more than100 papers per language), somewhat studied languages (with more than10 but fewer than 100 papers), and understudied / not studied languages (with fewer than 10 papers, several with not a single paper that we could find; note that for some of these, there exist past EEG/MEG studies). This table is not meant to serve as a comprehensive literature review, but to highlight the fact that for many, especially non-‘dominant’, languages, no fMRI investigations have been conducted, and if they have been, they tend to be clinical in nature (e.g., developing tools for pre-surgical mapping), to use the language as a control condition, and/or to be published in lower-impact journals.

| A characterization of the languages included in the current study—as well studied, somewhat studied, or understudied/not studied—with respect to past fMRI work. |  |
| --- | --- |
| i. Well Studied Languages (>100 papers per language) |  |
| Language | Sample Citation |
| Dutch | Snijders, T. M., Vosse, T., Kempen, G., Van Berkum, J. J., Petersson, K. M., & Hagoort, P. (2009). Retrieval and unification of syntactic structure in sentence comprehension: an fMRI study using word-category ambiguity. *Cerebral Cortex*, 19(7), 1493-1503. |
| English | Fedorenko, E., Behr, M. K., & Kanwisher, N. (2011). Functional specificity for high-level linguistic processing in the human brain. *Proceedings of the National Academy of Sciences*, 108(39), 16428-16433. |
| French | Pallier, C., Devauchelle, A. D., & Dehaene, S. (2011). Cortical representation of the constituent structure of sentences. *Proceedings of the National Academy of Sciences*, 108(6), 2522-2527. |
| German | Friederici, A. D., Meyer, M., & Von Cramon, D. Y. (2000). Auditory language comprehension: an event-related fMRI study on the processing of syntactic and lexical information. *Brain and Language*, 74(2), 289-300. |
| Japanese | Kim, J., Koizumi, M., Ikuta, N., Fukumitsu, Y., Kimura, N., Iwata, K., Watanabe, J., Yokoyama, S., Sato, S., Horie, K., & Kawashima, R. (2009). Scrambling effects on the processing of Japanese sentences: An fMRI study. *Journal of Neurolinguistics*, 22(2), 151-166. |
| Korean | Pallier, C., Dehaene, S., Poline, J. B., LeBihan, D., Argenti, A. M., Dupoux, E., & Mehler, J. (2003). Brain imaging of language plasticity in adopted adults: Can a second language replace the first? *Cerebral Cortex*, 13(2), 155-161. |
| Mandarin | Chee, M. W., Caplan, D., Soon, C. S., Sriram, N., Tan, E. W., Thiel, T., & Weekes, B. (1999). Processing of visually presented sentences in Mandarin and English studied with fMRI. *Neuron*, 23(1), 127-137. |
| Spanish | Brignoni-Perez, E., Jamal, N. I., & Eden, G. F. (2020). An fMRI study of English and Spanish word reading in bilingual adults. *Brain and Language*, 202, 104725. |
| ii. Somewhat Studied Languages (>10 but <100 papers) |  |
| Arabic | Mohtasib, R. S., Alghamdi, J. S., Baz, S. M., Aljoudi, H. F., Masawi, A. M., & Jobeir, A. A. (2021). Developing fMRI protocol for clinical use Comparison of 6 Arabic paradigms for brain language mapping in native Arabic speakers. *Neurosciences Journal*, 26(1), 45-55. |
| Basque | Quiñones, I., Amoruso, L., Pomposo Gastelu, I. C., Gil-Robles, S., & Carreiras, M. (2021). What can glioma patients teach us about language (re)organization in the bilingual brain: Evidence from fMRI and MEG. *Cancers*, 13(11), 2593. |
| Catalan | Perani, D., Abutalebi, J., Paulesu, E., Brambati, S., Scifo, P., Cappa, S. F., & Fazio, F. (2003). The role of age of acquisition and language usage in early, high-proficient bilinguals: An fMRI study during verbal fluency. *Human Brain Mapping*, 19(3), 170-182. |
| Danish | Buchweitz, A., Mason, R. A., Tomitch, L., & Just, M. A. (2009). Brain activation for reading and listening comprehension: An fMRI study of modality effects and individual differences in language comprehension. *Psychology and Neuroscience*, 2(2), 111-123. |
| Finnish | Hugdahl, K., Thomsen, T., Ersland, L., Rimol, L. M., & Niemi, J. (2003). The effects of attention on speech perception: an fMRI study. *Brain and Language*, 85(1), 37-48. |
| Greek | Kokkinos, V., Selviaridis, P., & Seimenis, I. (2021). Feasibility, contrast sensitivity and network specificity of language fMRI in presurgical evaluation for epilepsy and brain tumor surgery. *Brain Topography*, 34(4), 511-524. |
| Hebrew | Bitan, T., Kaftory, A., Meiri-Leib, A., Eviatar, Z., & Peleg, O. (2017). Phonological ambiguity modulates resolution of semantic ambiguity during reading: An fMRI study of Hebrew. *Neuropsychology*, 31(7), 759. |
| Hindi | Kumar, U., Padakannaya, P., Mishra, R. K., & Khetrapal, C. L. (2013). Distinctive neural signatures for negative sentences in Hindi: an fMRI study. *Brain Imaging and Behavior*, 7(2), 91-101. |
| Italian | Carota, F., Bozic, M., & Marslen-Wilson, W. (2016). Decompositional representation of morphological complexity: Multivariate fMRI evidence from Italian. *Journal of Cognitive Neuroscience*, 28(12), 1878-1896. |
| Norwegian | Lehtonen, M. H., Laine, M., Niemi, J., Thomsen, T., Vorobyev, V. A., & Hugdahl, K. (2005). Brain correlates of sentence translation in Finnish–Norwegian bilinguals. *NeuroReport*, 16(6), 607-610. |
| Polish | Bozic, M., Szlachta, Z., & Marslen-Wilson, W. D. (2013). Cross-linguistic parallels in processing derivational morphology: Evidence from Polish. *Brain and Language*, 127(3), 533-538. |
| Portuguese | Buchweitz, A., Mason, R. A., Tomitch, L., & Just, M. A. (2009). Brain activation for reading and listening comprehension: An fMRI study of modality effects and individual differences in language comprehension. *Psychology and Neuroscience*, 2(2), 111-123. |
| Russian | Axelrod, V., Bar, M., Rees, G., & Yovel, G. (2015). Neural correlates of subliminal language processing. *Cerebral Cortex*, 25(8), 2160-2169. |
| Swedish | Ettinger-Veenstra, V., McAllister, A., Lundberg, P., Karlsson, T., & Engström, M. (2016). Higher language ability is related to angular gyrus activation increase during semantic processing, independent of sentence incongruency. *Frontiers in Human Neuroscience*, 10, 110. |
| **iii. Understudied/Not Studied Languages (<10 papers)** |  |
| Afrikaans | Benjamin, C.F., Dhingra, I., Li, A.X., Blumenfeld, H., Alkawadri, R., Bickel, S., Helmstaedter, C., Meletti, S., Bronen, R.A., Warfield, S.K., & Spencer, D. D. (2018). Presurgical language fMRI: Current technical practices in epilepsy surgical planning. *bioRxiv*, 279117. |
| Armenian | No studies found. |
| Belarussian | No studies found. |
| Bulgarian | Kaiser, A., Kuenzli, E., Zappatore, D., & Nitsch, C. (2007). On females' lateral and males' bilateral activation during language production: a fMRI study. *International Journal of Psychophysiology*, 63(2), 192-198. |
| Czech | Brázdil, M., Chlebus, P., Mikl, M., Pažourková, M., Krupa, P., & Rektor, I. (2005). Reorganization of language-related neuronal networks in patients with left temporal lobe epilepsy—an fMRI study. *European Journal of Neurology*, 12(4), 268-275. |
| Farsi | Dehghani, M., Boghrati, R., Man, K., Hoover, J., Gimbel, S.I., Vaswani, A., Zevin, J.D., Immordino-Yang, M.H., Gordon, A.S., Damasio, A., & Kaplan, J. T. (2017). Decoding the neural representation of story meanings across languages. *Human Brain Mapping*, 38(12), 6096-6106. |
| Gujarati | Gupta, S. S. (2014). fMRI for mapping language networks in neurosurgical cases. *The Indian Journal of Radiology & Imaging*, 24(1), 37. |
| Hungarian | Kiss, M., Rudas, G., & Kozak, L. R. (2016, March). The outcome of fMRI language mapping is affected by patient fatigue. *European Congress of Radiology-ECR 2016*. |
| Irish | No studies found. |
| Latvian | No studies found. |
| Lithuanian | No studies found. |
| Marathi | No studies found. |
| Nepali | Mu, J., Xie, P., Yang, Z. S., Lu, F. J., Li, Y., & Luo, T. Y. (2006). Functional magnetic resonance image study on the brain areas involved in reading Chinese, English, and Nepali in Nepalese. *Zhong nan da xue xue bao. Yi xue ban. (Journal of Central South University. Medical Sciences.)*, 31(5), 759-762. |
| Romanian | No studies found. |
| Serbocroatian | Progovac, L., Rakhlin, N., Angell, W., Liddane, R., Tang, L., & Ofen, N. (2018). Diversity of grammars and their diverging evolutionary and processing paths: evidence from functional MRI study of Serbian. *Frontiers in Psychology*, 9, 278. |
| Slovene | Benjamin, C.F., Dhingra, I., Li, A.X., Blumenfeld, H., Alkawadri, R., Bickel, S., Helmstaedter, C., Meletti, S., Bronen, R.A., Warfield, S.K., & Spencer, D. D. (2018). Presurgical language fMRI: Current technical practices in epilepsy surgical planning. *bioRxiv*, 279117. |
| Swahili | No studies found. |
| Tagalog | No studies found. |
| Telugu | Agrawal, A., Hari, K. V. S., & Arun, S. P. (2018). How does reading expertise influence letter representations in the brain? An fMRI study. *Journal of Vision*, 18(10), 1161-1161. |
| Tamil | No studies found. |
| Turkish | No studies found. |
| Ukranian | No studies found. |
| Vietnamese | No studies found. |

**Supp. Table 2:** Results of linear mixed effects models. The analyses reported in the main text were supplemented with linear mixed effects models to ensure the robustness of the results to analytic procedure. These models also enabled us to examine inter-individual and inter-language/language-family variance (see also Supp. Figure 16). The key neural measures were predicted by a model that included a fixed effect of condition (specified below for each measure) and random intercepts by participant (n=86), language (n=45), language family (n=12), and fROI (n=6).

|  | Condition Sig | Participant | Language | Lang Family | ROI |
| --- | --- | --- | --- | --- | --- |
| **Response Strength Measures** |  |  |  |  |  |
| *EffectSize ~ Condition + (1 \| Participant) + (1 \| Language) + (1 \| Lang. Family) + (1 \| fROI)* |  |  |  |  |  |
| *Native-language > Degraded-language.* | p<0.001 | 0.42 | <0.01 | 0.01 | 0.40 |
| *Native-language > Unfamiliar-language* | p<0.001 | 0.37 | <0.01 | <0.01 | 0.34 |
| *Native-language > Spatial Working Memory (Hard).* | p<0.001 | 0.16 | <0.01 | 0.02 | 0.48 |
| *Native-language > Math (Hard).* | p<0.001 | 0.14 | <0.01 | 0.02 | 0.25 |
| **Lateralization measures (response strength and activation extent):** |  |  |  |  |  |
| *EffectSize ~ Hemisphere + (1 \| Participant) (1 \| Language) + (1 \| Lang. Family) + (1 \| fROI)* |  |  |  |  |  |
| *Left Hemisphere > Right Hemisphere Strength* | p<0.01 | 0.33 | 0.04 | 0.01 | 0.77 |
| *Supra-threshold voxels in left > Right hemi* | p<0.01 | 11,701 | 2,383 | 0.00 | 91,579 |
| **Lateralization measures (functional correlations):** |  |  |  |  |  |
| *EffectSize ~ Hemisphere + (1 \| Participant) + (1 \| Language) + (1 \| Lang. Family)* |  |  |  |  |  |
| *Story Comprehension Left > Right* | p<0.01 | 0.02 | <0.01 | <0.01 | NA |
| *Resting State Left > Right Hemisphere* | p<0.01 | 0.01 | <0.01 | <0.01 | NA |
| **Within- and between-network correlation measures.** |  |  |  |  |  |
| Here, networks (either language-language (pairs of fROIs within the language network) or language-MD (pairs of fROIs straddling network boundaries)) were modeled as a fixed effects: |  |  |  |  |  |
| *EffectSize ~ Systems + (1 \| Language) + (1 \| Subject) + (1 \| Language Family)* |  |  |  |  |  |
| *Story comprehension Within language network > lang-MD correlations* | p<0.01 | <0.01 | <0.01 | <0.01 | NA |
| *Resting state Within language network > lang-MD correlations* | p<0.01 | <0.01 | <0.01 | <0.01 | NA |

**Supplementary Table 3:** Information on the language background of all participants. Participants are numbered 1-86 in column 1 (the number in parentheses is the UID (unique ID)— the internal lab identifier that is used in all the data tables and files on OSF: https://osf.io/cw89s/.). For each language listed in columns 2-4, we report in parentheses i) age of acquisition, ii) self-reported spoken proficiency (the average of self-reported spoken comprehension proficiency and speaking proficiency) on a scale from 1 (very basic proficiency) to 5 (native-like proficiency), iii) self-reported written proficiency (the average of self-reported written comprehension proficiency and writing proficiency) on the same 1-5 scale, and iv) environment in which the language was acquired (‘home’ indicates that one or both parents speak the language, ‘class’ indicates a formal language class either in high school or university). Listed under ‘Native language(s)’ is/are the language(s) that the participant listed as having learnt before the age of 6, with one or both parents speaking the language. Listed under ‘Language(s) spoken fluently’ is/are the language(s) with a self-reported spoken proficiency of 3 and above. Listed under ‘Language(s) with some familiarity’ is/are the rest of the languages reported by the participant.

| Participant Number | Native language(s) | Language(s) spoken fluently | Language(s) with some familiarity |
| --- | --- | --- | --- |
| 1 (544) | Arabic (0; 5; 5; home/class) | English (4; 5; 5; class) | French (15; 3; 3.5; class)<br>German (25; 2; 3; class) |
| 2 (561) | Arabic (0; 5; 5; home/class) | English (5; 5; 5; class) | French (6; 2; 2; class)<br>German (19; 3.5; 3.5; class)<br>Spanish (19; 3.5; 3.5; class)<br>Italian (19; 2; 2; class) |
| 3 (182) | Hebrew (0; 5; 5; home/class) | English (8; 5; 5; class) | French (25; 2; 2; class)<br>Arabic (14; 1; 1.5; class)<br>American Sign Language (30; 2; 1; class) |
| 4 (506) | Hebrew (0; 5; 5; home/class)<br>English (0; 5; 5; home/class) | Spanish (7; 4; 4; class) | German (21; 2; 2; class)<br>Swedish (21; 2; 2; class) |
| 5 (458) | Vietnamese (0;5; 5; home/class) | English (10; 3.5; 4; class) |  |
| 6 (570) | Vietnamese (0;5;4.5; home/class) | English (7; 5; 5; home/class) | French (13; 2.5; 2.5; home/class)<br>Spanish (13; 1.5; 2; home/class) |
| 7 (580) | Tagalog (0; 5; 5; home)<br>English (0; 5; 5; home/class) |  |  |
| 8 (467) | Tamil (1; 5; 5; home/class)<br>English (3; 5; 5; home/class) |  | Japanese (21; 3.5; 4; class)<br>German (18; 2; 2; class) |
| 9 (500) | Tamil (1; 5; 5; home)<br>English (3; 5; 5; home/class) |  |  |
| 10 (800) | Telugu (0; 5; 1; home)<br>English (1; 5; 5; home/class) |  | Hindi (1; 5; 4; home/class)<br>French (14; 2; 2; class)<br>Gujarati (4; 3; 1; home/class) |
| 11 (451) | Afrikaans (0; 5; 5; home/class)<br>English (2; 5; 5; home/class) |  | Dutch (9; 2; 2.5; class)<br>German (16; 2; 2; class) |
| 12 (455) | Afrikaans (0; 5; 5; home/class) | English (5; 4.5; 5; class) | Greek (20; 1.5; 1.5; class)<br>Hebrew (20; 1.5; 1.5; class) |
| 13 (454) | Armenian (0; 5; 5; home/class) | Russian (6; 5; 4.5; class)<br>English (9; 5; 5; home/ class) | French (21; 2; 2; class) |
| 14 (493) | Armenian (0; 5; 4; home/class) | English (10; 5; 5; home/class)<br>Russian (0; 4; 4; class) | French (15; 2; 1.5; class) |
| 15 (543) | Belarusian (5; 4; 4; home/class)<br>Russian (0; 5; 5; home/class) | English (10; 4; 4.5; home/class) | German (19; 2; 2; class) |
| 16 (611) | Belarusian (1; 5; 5; home/class)<br>Russian (0; 5; 5; home/class)<br>English (3; 4.5; 4.5; home/class) |  | Lithuanian (11; 3.5; 3.5; home/ class)<br>French (18; 3.5; 3; home/ class)<br>Polish (10; 3; 3; home/ class)<br>German (12; 2.5; 2.5; class)<br>Latvian (2; 2.5; 2.5; class)<br>Georgian (24; 2; 1.5; class)<br>Old Church Slavic (21; 1; 2.5; class)<br>Latin (18; 1; 2; class)<br>Sanskrit (21; 1; 2; class) |
| 17 (513) | Bulgarian (0; 5; 5; home/class)<br>English (3; 5; 5; home/ class) | Spanish (21; 4; 4; home/ class) | Russian (6; 3.5; 3.5; class)<br>French (14; 3; 3; class)<br>German (19; 2; 2; class) |
| 18 (517) | Bulgarian (0; 5; 5; home/class)<br>Russian (0; 5; 5; home/class)<br>Spanish (0; 5; 5; home/class)<br>German (0; 5; 5; home/class) | English (10; 4.5; 5; class) |  |
| 19 (450) | Catalan (0; 5; 5; home/class)<br>Spanish (0; 5; 5; home/class) | English (4; 5; 5; class) | Serbo-croatian (23; 1; 1; class) |
| 20 (464) | Catalan (0; 5; 5; home/class)<br>Spanish (0; 5; 5; home/class) | English (0; 5; 5; class) | German (13; 2; 2; class) |
| 21 (638) | Czech (details missing) | English (15; 4.5; 4.5; class) | German (8; 2; 3; class)<br>Russian (17; 2; 1.5; class)<br>Lithuanian (23; 1.5; 1.5; class) |
| 22 (647) | Czech (0; 5; 5; home/class) | English (14; 5; 5; class) | Russian (6; 3.5; 3.5; class)<br>Spanish (42; 3; 3; class)<br>German (0; 3; 3; class)<br>French (42; 2; 2; class) |
| 23 (507) | Danish (0; 5; 5; home/class)<br>English (0; 5; 5; home/class) | French (6; 5; 3.5; home/class)<br>Spanish (15; 4; 4; home/class) | Norwegian (10; 2.5; 2.5; home/class)<br>Swedish (19; 2.5; 2.5; home)<br>German (12; 2; 1.5; home/class) |
| 24 (508) | Danish (0; 5; 5; home/class) | English (7; 5; 5; home/class) | Norwegian (11; 2; 3; class)<br>Swedish (11; 2.5; 1.5; class)<br>German (7; 2; 1.5; class)<br>French (15; 1.5; 1; class)<br>Italian (20; 1; 1; class) |
| 25 (463) | Dutch (0; 5; 5; home/class) | English (10; 4.5; 4.5; class)<br>Portuguese (15; 4.5; 4.5; class) | French (0; 5; 5; class)<br>German (0; 5; 5; class) |
| 26 (481) | Dutch (0; 5; 5; home/class)<br>English (0; 5; 5; home/class) |  | German (0; 2.5; 3; home/class)<br>French (12; 1.5; 1.5; class)<br>Spanish (18; 1.5; 1.5; class) |
| 27 (492) | English (0; 5; 5; home/class) | Spanish (13; 2; 2; class) |  |
| 28 (502) | English (0; 5; 5; home/class) |  | German (11; 2.5; 2; class)<br>French (10; 1; 1; class)<br>Latin (4; 1; 1; class) |
| 29 (443) | Farsi (0; 5; 5; home/class) | English (10; 5; 5; home) | German (4; 3.5; 3.5; home/class)<br>Spanish (18; 2; 2; class)<br>Turkish (2; 2; 1; class)<br>Greek (27; 1.5; 2; class)<br>Arabic (12; 1.5; 2; class)<br>Hungarian (2; 2; 1; class) |
| 30 (617) | Farsi (0; 5; 5; home/class) | English (details missing) |  |
| 31 (462) | French (0; 5; 5; home/class)<br>English (0; 5; 5; home/class) |  | German (0; 5; 5; home/class) |
| 32 (480) | French (0; 5; 5; home/class) | English (7; 5; 5; class) | Spanish (12; 1.5; 3; class)<br>German (23; 1.5; 2; class) |
| 33 (457) | German (0; 5; 5; home/class) | English (7; 5; 5; home/class) | Mandarin (0; 3; 2; home/class)<br>Latin (10; 1; 1; class) |
| 34 (482) | German (0; 5; 5; home/class)<br>Romanian (3; 5; 4.5; home) | English (11; 4; 4; home/class) | Hungarian (3; 2; 2; home) |
| 35 (496) | Greek (0; 5; 5; home/class) | English (8; 5; 4.5; home/class) | German (6; 3; 3.5; class) |
| 36 (548) | Greek (0; 5; 5; home/class) | English (7; 4; 4; home/class) | French (11; 2.5; 3; class)<br>German (14; 2; 2; class)<br>Portuguese (17; 2; 2; class)<br>Italian (24; 1; 2; class) |
| 37 (799) | Gujarati (0; 4.5; 2; home/class)<br>English (0; 5; 5; home/class) |  | Hindi (5; 4.5; 3; home/class)<br>Arabic (19; 2; 2; class)<br>Bengali (20; 2; 2; class)<br>Latin (11; 2; 2; class) |
| 38 (808) | Gujarati (0; 4; 3; home)<br>Italian (0; 5; 5; home) | English (11; 5; 5; class) | French (3; 5; 5; home/class)<br>Spanish (14; 4; 4; class) |
| 39 (470) | Hindi (0; 5; 5; home/class) | English (4; 5; 5; home/class) |  |
| 40 (504) | Hindi (2; 5; 5; home/class) | English (5; 5; 5; home/class) | Marathi (5; 3.5; 3.5; home/class)<br>Marwadi (NA; 3; 3; class) |
| 41 (618) | Irish (1; 5; 5; home/class)<br>English (1; 4; 4; home/class) |  |  |
| 42 (620) | Irish (0; 5; 4; home/class)<br>English (0; 5; 5; home/class) |  | French (12; 3; 3; home/class)<br>German (10; 2; 2; class) |
| 43 (437) | Italian (0; 5; 5; home) | English (0; 5; 5; class) | Chinese (23; 1; 1; class) |
| 44 (444) | Italian (0; 5; 5; home/class) | English (11; 4; 4; class) | French (11; 2; 3; class)<br>Spanish (29; 2; 2.5; class)<br>German (27; 1; 1.5; class) |
| 45 (634) | Latvian (0; 5; 5; home/class)<br>English (0; 5; 5; home/class) |  | French (4; 3; 4; class)<br>Italian (0; 1; 1; class) |
| 46 (635) | Latvian (0; 5; 5; home/class) | English (17; 5; 4; class) | Russian (7; 3.5; 2; class) |
| 47 (565) | Lithuanian (0; 5; 5; home/class) | English (4; 5; 5; class)<br>French (4; 5; 4.5; class) | German (11; 4; 3.5; class)<br>Spanish (13; 3.5; 3.5; class) |
| 48 (579) | Lithuanian (0; 5; 5; home/class) | English (6; 5; 5; home/class) | Spanish (19; 3; 2.5; class)<br>Russian (10; 2; 1; class) |
| 49 (810) | Marathi (0; 5; 5; home/class)<br>English (2; 5; 5; home/class)<br>Hindi (2; 5; 5; home/class) |  | French (13; 1.5; 1; class) |
| 50 (813) | Marathi (0; 5; 5; home/class) | English (3; 5; 5; class) | Urdu (1; 3; 1; home/class)<br>Punjabi (10; 2.5; 1; class)<br>French (25; 1; 1; class) |
| 51 (515) | Nepali (0; 5; 3; home/class)<br>English (2; 5; 5; home/class) | Hindi (0; 5; 3; class) |  |
| 52 (581) | Nepali (0; 5; 5; home/class)<br>English (2; 5; 5; home/class) |  | Hindi (8; 4.5; 2.5; other) |
| 53 (460) | Norwegian (0; 5; 5; home/class) | English (7; 4.5; 4.5; class) | Swedish (5; 3.5; 4; class)<br>Danish (10; 2.5; 3.5; class)<br>Spanish (13; 2; 2.5; class) |
| 54 (469) | Norwegian (0; 5; 5; home)<br>Swedish (0; 4.5; 4; home) | English (10; 5; 5; class) | Spanish (13; 2; 2; class) |
| 55 (446) | Polish (0; 5; 5; home/class) | English (5; 5; 5; home/class)<br>German (15; 4; 4; class)<br>Slovene (19; 4; 4; class)<br>Serbocroatian (21; 4; 4; class) | Lithuanian (23; 3; 4; class)<br>Russian (21; 3; 4; class)<br>Bulgarian (26; 3; 3; class)<br>Albanian (28; 3; 3; class)<br>Latvian (25; 2; 3; class)<br>Czech (27; 3; 3; class)<br>Ukrainian (25; 2; 2.5; class)<br>Upper Sorbian (26; 2; 2.5; class)<br>French (24; 2; 2.5; class)<br>Norwegian (23; 1.5; 1.5; class)<br>Macedonian (29; 2; 2.5; class) |
| 56 (445) | Polish (0; 5; 5; home/class)<br>English (0; 4; 4; home/class) | Serbocroatian (18; 4; 4; class) | Slovene (20; 3.5; 3.5; class)<br>German (20; 3.5; 3.5; home/class)<br>French (25; 1.5; 2.5; class) |
| 57 (459) | Portuguese (0; 5; 5; home/class) | English (0; 4.5; 5; class)<br>Spanish (0; 4.5; 4.5; home/class) | French (10; 2; 2; class)<br>Mandarin (16; 3; 3; class) |
| 58 (484) | Portuguese (0; 5; 5; home/class) | English (10; 4; 4; class) | Spanish (13; 3; 2; class)<br>French (25; 2; 1; class)<br>Russian (28; 1; 1; class) |
| 59 (501) | Romanian (0; 5; 5; home) | English (10; 4.5; 4; class) | French (12; 1.5; 1.5; class) |
| 60 (509) | Romanian (0; 5; 5; home/class) | English (6; 5; 5; class) | French (11; 2; 2.5; class) |
| 61 (440) | Russian (0; 5; 5; home/class) | English (10; 3; 3; class) |  |
| 62 (538) | Russian (0; 5; 5; home/class) | English (7; 5; 4.5; home/class) | German (18; 3; 2.5; home/class) |
| 63 (468) | Serbocroatian (0; 5; 5; home/class) | English (5; 5; 5; class) | Slovene (5; 3.5; 3.5; class)<br>Italian (8; 3; 2.5; class)<br>Spanish (8; 3; 2; class)<br>German (10; 2; 2; class) |
| 64 (546) | Serbocroatian(0; 5; 5; home/class) | English (4; 5; 5; class) | Italian (15; 3; 3; class)<br>German (15; 2; 2; class) |
| 65 (503) | Slovene (0; 5; 5; home/class) | English (14; 3; 3.5; class) | Serbocroatian (1; 5; 5; home/class)<br>German (8; 2; 1; class)<br>Italian (2; 2; 1; home/class)<br>Dutch (24; 1.5; 1; class) |
| 66 (497) | Spanish (0; 5; 5; home/class)<br>English (3; 4; 4; home/class) |  | French (19; 3; 3; class)<br>German (4; 3; 3; class) |
| 67 (547) | Spanish (0; 5; 5; home/class)<br>English (5; 3.5; 3.5; home/class) |  |  |
| 68 (514) | Swedish (0; 5; 5; home/class) | English (9; 5; 5; home/class) | Spanish (0; 4; 4; class)<br>Mandarin (31; 1; 1; class) |
| 69 (542) | Swedish (0; 5; 5; home/class) | English (1; 5; 5; home/class) | French (11; 3; 2.5; class)<br>Mandarin (19; 2; 2; class)<br>Spanish (24; 1.5; 1; class) |
| 70 (490) | Ukrainian (0; 4.5; 4.5; home)<br>Russian (0; 5; 5; home) | English (6; 5; 5; class) | French (9; 2; 2; class) |
| 71 (495) | Ukrainian (0; 5; 5; home/class)<br>Russian (0; 5; 5; home/class)<br>English (3; 4; 4; home/class) |  | German (11; 1.5; 1.5; class) |
| 72 (628) | Basque (0; 5; 5; home/class)<br>Spanish (0; 5; 5; home/class) | English (11; 4; 4; home/class) | French (13; 3; 2.5; class) |
| 73 (809) | Basque (2; 5; 5; home/class)<br>Spanish (0; 5; 5; home/class) | English (5; 5; 5; class) | Italian (24; 4; 4; class)<br>French (12; 2; 2; class)<br>Portuguese (24; 3.5; 3.5; class) |
| 74 (465) | Japanese (0; 5; 5; home/class) | English (12; 5; 5; class) | German (18; 2; 2; class) |
| 75 (512) | Japanese (1; 5; 4; home/class) | English (10; 5; 5; home/class) | Mandarin (1; 3.5; 2.5; class) |
| 76 (466) | Korean (1; 5; 5; home/class) | English (10; 3.5; 4; class) |  |
| 77 (488) | Korean (0; 4.5; 3.5; home/class) | English (13; 4; 4; class) |  |
| 78 (804) | Swahili (0; 5; 5; home/class)<br>English (0; 5; 5; home/class)<br>Kimeru (0; 5; 1.5; home) |  |  |
| 79 (478) | Mandarin (0; 4; 3; home/class)<br>English (3; 5; 5; home/class) |  | Japanese (15; 2; 2; class) |
| 80 (491) | Mandarin (0; 5; 5; home/class) | English (7; 4.5; 4.5; home/class) | Japanese (21; 2; 2; class) |
| 81 (510) | Turkish (0; 5; 5; home/class) | English (10; 4.5; 5; class) | German (11; 4; 4; class)<br>Spanish (19; 3; 3; class) |
| 82 (533) | Turkish (0; 5; 5; home/class) | Arabic (15; 5; 5; class)<br>English (15; 5; 5; class) | French (28; 2; 3; class)<br>Farsi (32; 2; 2; class) |
| 83 (619) | Finnish (0; 5; 5; home/class)<br>Swedish (0; 5; 5; home/class) | English (20; 4; 4; class) |  |
| 84 (648) | Finnish (0; 5; 5; home/class) | English (9; 4; 5; class) | Swedish (4; 3; 3.5; class)<br>German (14; 3; 3; class) |
| 85 (471) | Hungarian (0; 5; 5; home)<br>English (3; 5; 5; home) | German (7; 5; 5; home) | French (12; 3; 3; class)<br>Spanish (21; 2; 2.5; class) |
| 86 (518) | Hungarian (0; 5; 5; home/class) | English (8; 4; 4.5; class) | German (22; 2.5; 2; class)<br>Spanish (24; 2; 2.5; class)<br>Latin (14; 1; 2; class) |

**Supplementary Table 4:** Information on the gender and age of the participants (at testing), as well as the number of participants tested per language. The table is sorted alphabetically by language family, and then by language.

| **Native Language** | **Language Family** | **Number of Participants** | **Participant Sex and Age** |
| --- | --- | --- | --- |
| Arabic | Afro-Asiatic | 2 | Male (28), Female (27) |
| Hebrew | Afro-Asiatic | 2 | Male (31), Female (26) |
| Swahili | Atlantic-Congo | 1 | Female (19) |
| Tagalog | Austronesian | 1 | Male (22) |
| Vietnamese | Austroasiatic | 2 | Male (21), Female (20) |
| Tamil | Dravidian | 2 | Male (25), Female (22) |
| Telugu | Dravidian | 1 | Male (28) |
| Afrikaans | Indo-European | 2 | Male (37), Female (25) |
| Armenian | Indo-European | 2 | Male (23), Female (30) |
| Belarusian | Indo-European | 2 | Male (23), Female (27) |
| Bulgarian | Indo-European | 2 | Male (37), Female (36) |
| Catalan | Indo-European | 2 | Male (25), Female (27) |
| Czech | Indo-European | 2 | Male (44), Female (27) |
| Danish | Indo-European | 2 | Male (32), Female (26) |
| Dutch | Indo-European | 2 | Male (32), Female (25) |
| English | Indo-European | 2 | Male (23), Female (25) |
| Farsi | Indo-European | 2 | Male (30), Female (32) |
| French | Indo-European | 2 | Male (29), Female (25) |
| German | Indo-European | 2 | Male (23), Female (30) |
| Greek | Indo-European | 2 | Male (26), Female (25) |
| Gujarati | Indo-European | 2 | Male (27), Female (27) |
| Hindi | Indo-European | 2 | Male (27), Female (22) |
| Irish | Indo-European | 2 | Male (26), Female (30) |
| Italian | Indo-European | 2 | Male (29), Female (29) |
| Latvian | Indo-European | 2 | Male (45), Female (25) |
| Lithuanian | Indo-European | 2 | Male (22), Female (19) |
| Marathi | Indo-European | 2 | Male (31), Female (28) |
| Nepali | Indo-European | 2 | Male (21), Female (24) |
| Norwegian | Indo-European | 2 | Male (22), Female (25) |
| Polish | Indo-European | 2 | Male (31), Female (31) |
| Portuguese | Indo-European | 2 | Male (19), Female (34) |
| Romanian | Indo-European | 2 | Male (19), Female (20) |
| Russian | Indo-European | 2 | Male (32), Female (23) |
| Serbocroatian | Indo-European | 2 | Male (28), Female (33) |
| Slovene | Indo-European | 1 | Female (24) |
| Spanish | Indo-European | 2 | Male (31), Female (41) |
| Swedish | Indo-European | 2 | Male (32), Female (31) |
| Ukrainian | Indo-European | 2 | Male (20), Female (19) |
| Basque | Isolate | 2 | Male (28), Female (25) |
| Japanese | Japonic | 2 | Male (29), Female (19) |
| Korean | Koreanic | 2 | Male (31), Female (29) |
| Mandarin | Sino-Tibetan | 2 | Male (25), Female (20) |
| Turkish | Turkic | 2 | Male (33), Female (30) |
| Finnish | Uralic | 2 | Male (37), Female (34) |
| Hungarian | Uralic | 2 | Male (25), Female (30) |

